# TCDD-Induced m⁶A Profiling Identifies SUV39H2 as a Positive Regulator of Aryl Hydrocarbon Receptor Signaling

**DOI:** 10.64898/2026.11.19.767661

**Authors:** Gabin Kwon, HongDuck Yun, Visanu Wanchai, Stephen Safe, KyoungHyun Kim

## Abstract

The aryl hydrocarbon receptor (AHR) mediates cellular responses to environmental contaminants, including 2,3,7,8-tetrachlorodibenzo-*p*-dioxin (TCDD). Although AHR-dependent transcriptional regulation is well characterized, the contributions of RNA modifications and chromatin-associated regulators to AHR signaling remain incompletely understood. In this study, we investigated TCDD-associated changes in *N*⁶-methyladenosine (m⁶A) RNA modification and identified suppressor of variegation 3-9 homolog 2 (SUV39H2) as a previously unrecognized positive regulator of AHR signaling. Exploratory methylated RNA immunoprecipitation sequencing (MeRIP-seq) and matched RNA-seq analyses were performed in vehicle- and TCDD-treated HepG2 cells, followed by independent molecular and functional validation. TCDD increased m⁶A enrichment and mRNA stability of canonical AHR-responsive transcripts, including *CYP1A1*, *CYP1A2*, and *AHRR*, accompanied by increased gene expression. Notably, *SUV39H2* was identified as a TCDD-responsive m⁶A-enriched transcript exhibiting increased mRNA stability and protein expression without a substantial change in steady-state mRNA abundance. Functional studies demonstrated that SUV39H2 depletion and pharmacological inhibition attenuated TCDD-induced AHR target gene expression and transcriptional activity, whereas SUV39H2 overexpression enhanced AHR-dependent responses. CUT&RUN-qPCR analyses further implicated SUV39H2 in AHR-associated transcriptional regulation at the *CYP1A1* enhancer. Consistent with the cellular findings, pharmacological inhibition of SUV39H2 attenuated TCDD-induced expression of AHR-responsive genes in mouse liver. Collectively, these findings identify SUV39H2 as a previously unrecognized positive regulator of AHR signaling and suggest that TCDD-associated m⁶A modification and mRNA stabilization contribute to the regulation of AHR-responsive transcripts and chromatin-associated regulatory factors.

## Introduction

The aryl hydrocarbon receptor (AHR) is a ligand-activated transcription factor that mediates cellular responses to environmental contaminants and endogenous signaling molecules. Initially characterized as a receptor involved in the toxic effects of dioxins and related compounds, AHR is now recognized as an important regulator of xenobiotic metabolism, inflammation, cellular differentiation, and carcinogenesis (Denison and Nagy 2003; Safe et al. 2013, 2020; Kim 2024). Among its exogenous ligands, 2,3,7,8-tetrachlorodibenzo-*p*-dioxin (TCDD) is a potent and persistent AHR agonist widely used to investigate AHR-dependent signaling. Upon ligand activation, AHR translocates to the nucleus, heterodimerizes with the AHR nuclear translocator (ARNT), and binds to dioxin response elements (DREs) within regulatory regions of target genes. This process induces xenobiotic-metabolizing enzymes, including CYP1A1 and CYP1A2, as well as the AHR repressor (AHRR), which contributes to negative feedback regulation of AHR signaling (Mimura et al. 1999; Beischlag et al. 2008). Beyond its canonical role in xenobiotic metabolism, AHR exhibits ligand- and cell-specific transcriptional activities influenced by interactions with transcriptional coregulators and chromatin-associated proteins (Beischlag et al. 2008; Safe et al. 2020). However, the molecular mechanisms through which additional chromatin-associated regulators modulate AHR-dependent transcription remain incompletely understood.

Chromatin-associated proteins regulate transcription factor activity by modifying histones, altering chromatin organization, and interacting with transcriptional regulatory complexes. Suppressor of variegation 3-9 homolog 2 (SUV39H2), also known as KMT1B under the standardized nomenclature for chromatin-modifying enzymes (Allis et al. 2007), is a histone lysine methyltransferase primarily associated with histone H3 lysine 9 (H3K9) methylation, heterochromatin formation, and transcriptional repression. However, SUV39H2 may also function as a transcriptional coactivator in specific cellular contexts. Askew et al. (2017) demonstrated that SUV39H2 interacts with the androgen receptor (AR) and its transcriptional coregulator MAGE-A11, is recruited to an androgen-responsive enhancer, and enhances androgen-dependent transcriptional activity. These findings suggest that SUV39H2 can participate in ligand-dependent transcription through mechanisms extending beyond its conventional role in heterochromatin formation. Although AR and AHR belong to distinct transcription factor families, both depend on interactions with regulatory proteins to control target gene expression. Thus, SUV39H2 represents a potential regulator of AHR-dependent transcription. Nevertheless, whether SUV39H2 participates in AHR signaling or modulates transcriptional responses to environmental AHR ligands has not been established.

In addition to chromatin-mediated regulation, post-transcriptional mechanisms influence the abundance and stability of transcripts involved in cellular signaling. N⁶-methyladenosine (m⁶A) is one of the most prevalent internal modifications of eukaryotic mRNA and is frequently enriched near stop codons and within 3′ untranslated regions (UTRs) (Dominissini et al. 2012; Meyer et al. 2012). Transcriptome-wide profiling using methylated RNA immunoprecipitation sequencing (MeRIP-seq) has enabled the identification of m⁶A-enriched RNA regions and the characterization of differential methylation patterns under various biological conditions (Dominissini et al. 2013; Meng et al. 2014). m⁶A modification is regulated by methyltransferase complexes containing METTL3 and METTL14, demethylases such as FTO and ALKBH5 (Jia et al. 2011; Zheng et al. 2013), and reader proteins, including YTH-domain-containing proteins and IGF2BP family members, that influence RNA processing, stability, and translation (Wang et al. 2014, 2015; Xiao et al. 2016; Huang et al. 2018). Depending on the associated reader proteins and cellular context, m⁶A modification can promote mRNA degradation, enhance transcript stability, or regulate translation. Recent studies have implicated m⁶A regulation in cellular responses to environmental chemicals. Shen et al. (2024) demonstrated that naphtho[2,1-*a*]pyrene exposure enhances CYP1A1 mRNA stability through an m⁶A-dependent mechanism involving METTL14 and IGF2BP3. Furthermore, Nakano et al. (2025) reported that cigarette smoking modulates m⁶A-related regulatory mechanisms and affects CYP1A1 induction through regulation involving ARNT and AHRR. These findings suggest that RNA methylation contributes to AHR-responsive gene expression. However, whether TCDD-associated m⁶A changes influence the expression of chromatin-associated regulators, particularly SUV39H2, and thereby contribute to AHR-dependent transcription remains unknown.

In this study, we investigated TCDD-associated changes in m⁶A RNA modification to identify previously unrecognized regulators of AHR signaling, with particular emphasis on SUV39H2. Exploratory transcriptome-wide m⁶A profiling and matched RNA-seq analyses were performed in vehicle- and TCDD-treated HepG2 cells to identify candidate m⁶A-enriched transcripts. We examined TCDD-associated changes in m⁶A enrichment and mRNA stability of canonical AHR-responsive genes, including *CYP1A1*, *CYP1A2*, and *AHRR*, and further investigated SUV39H2 regulation following TCDD exposure. The functional contribution of SUV39H2 to AHR signaling was evaluated using genetic depletion, overexpression, pharmacological inhibition, reporter assays, and CUT&RUN-qPCR. In addition, the effects of SUV39H2 inhibition on TCDD-responsive gene expression were examined in mouse liver. Together, these studies identify SUV39H2 as a previously unrecognized positive regulator of AHR signaling and suggest a potential relationship between TCDD-associated m^6^A RNA methylation, transcript stability, and chromatin-associated regulation of AHR-dependent transcription.

## Materials and Methods

### Cell culture, reagents, and treatments

Human hepatocellular carcinoma HepG2 and Hep3B cells were obtained from the American Type Culture Collection (ATCC, Manassas, VA, USA). HepG2 cells were maintained in Dulbecco’s modified Eagle’s medium (DMEM) supplemented with 5% fetal bovine serum (FBS), whereas Hep3B cells were cultured in DMEM containing 10% FBS. Both cell lines were maintained in medium supplemented with 1% penicillin–streptomycin at 37 °C in a humidified atmosphere containing 5% CO₂. To activate aryl hydrocarbon receptor (AHR) signaling, cells were treated with 10 nM 2,3,7,8-tetrachlorodibenzo-p-dioxin (TCDD; AccuStandard, New Haven, CT, USA) dissolved in dimethyl sulfoxide (DMSO). Cells were exposed to TCDD for 2 h in selected experiments or for the indicated durations in time-course experiments (0, 8, and 20 h). For MeRIP-seq, cells were treated with 10 nM TCDD for 5 h. Vehicle-treated cells received an equivalent volume of DMSO. For pharmacological inhibition of SUV39H2, OTS186935 (MedChemExpress, Cat. No. HY-122182; CAS No. 2093401-33-1; purity 98.64%) was dissolved in DMSO. HepG2 and Hep3B cells were treated with OTS186935 at concentrations of 1 or 3 µM for 2 h, as indicated. Corresponding vehicle controls were included in each experiment.

### Methylated RNA immunoprecipitation sequencing (MeRIP-seq)

Methylated RNA immunoprecipitation sequencing (MeRIP-seq) was performed using an antibody-based enrichment approach adapted from established protocols (Dominissini et al. 2012, 2013; Meyer et al. 2012). HepG2 cells were treated with vehicle or 10 nM TCDD for 5 h. Total RNA was isolated using TRIzol reagent (Invitrogen, Carlsbad, CA, USA) according to the manufacturer’s instructions. RNA quantity and integrity were assessed using an Agilent 2100 Bioanalyzer and RNA 6000 Nano Kit (Agilent Technologies, Santa Clara, CA, USA). More than 50 µg of total RNA with an RNA integrity number (RIN) greater than 7.0 was subjected to poly(A)⁺ mRNA enrichment using oligo(dT)-conjugated magnetic beads (Invitrogen). Purified mRNA was fragmented into approximately 100-nucleotide fragments using divalent cations at elevated temperature. Fragmented RNA was incubated with an anti-m⁶A antibody (Cat. No. 202003; Synaptic Systems, Germany) for 2 h at 4 °C in immunoprecipitation (IP) buffer containing 50 mM Tris-HCl, 750 mM NaCl, 0.5% IGEPAL CA-630, and 0.5 µg/µL bovine serum albumin. Antibody-bound RNA fragments were captured using protein A beads and eluted with IP buffer supplemented with 6.7 mM free m⁶A. The eluted RNA was recovered by ethanol precipitation. Immunoprecipitated RNA and corresponding input RNA were used to construct strand-specific cDNA libraries using a dUTP-based method. The average library insert size was approximately 100 ± 50 bp. Paired-end sequencing (2 × 150 bp) was performed on an Illumina NovaSeq 6000 platform by LC Sciences (Houston, TX, USA). One biological sample per treatment condition was analyzed, generating four libraries: vehicle input, vehicle m⁶A-IP, TCDD input, and TCDD m⁶A-IP. The sequencing experiment was designed as an exploratory transcriptome-wide analysis to identify candidate TCDD-responsive m⁶A-enriched transcripts for subsequent experimental validation.

### Bioinformatics analysis

Raw sequencing reads were processed using Cutadapt (Martin 2011) and custom Perl scripts to remove adapter sequences, low-quality reads, and reads containing undetermined bases. Read quality was assessed using FastQC (Andrews 2010), and high-quality reads were aligned to the human reference genome (GRCh38/hg38; Ensembl release 107) using Bowtie2 (Langmead and Salzberg 2012).

m⁶A-enriched regions and differential m⁶A peaks between vehicle- and TCDD-treated HepG2 cells were identified using exomePeak2 (Zhou et al. 2026), with matched input RNA libraries used to account for differences in transcript abundance. Candidate differential peaks were evaluated based on log₂ fold changes, nominal *P* values, and false discovery rate (FDR)-adjusted *P* values. Peaks were annotated using ChIPseeker (Yu et al. 2015), sequence motifs were analyzed using MEME and HOMER (Heinz et al. 2010; Bailey et al. 2015), and enrichment patterns were visualized using the Integrative Genomics Viewer (IGV) (Robinson et al. 2011). Transcript abundance was estimated from input RNA-seq data using StringTie (Pertea et al. 2015) and expressed as fragments per kilobase of transcript per million mapped reads (FPKM).

Differential gene expression was analyzed using edgeR (Robinson et al. 2010). Because only one biological sample was available per treatment condition, biological dispersion could not be estimated directly. Therefore, a fixed dispersion of 0.04, corresponding to a biological coefficient of variation of 20%, was used for exploratory analysis, following the approach described in the edgeR User’s Guide for RNA-seq experiments without biological replicates (Chen et al. 2025). This assumed dispersion enabled preliminary identification of candidate genes but did not replace an experimentally estimated biological variance. Genes with an absolute log₂ fold change greater than 1 and a nominal *P* value below 0.05 were considered candidate differentially expressed genes. Kyoto Encyclopedia of Genes and Genomes (KEGG) pathway enrichment was assessed using a hypergeometric test, with a nominal *P* value below 0.05 (Kanehisa and Goto 2000).

Publicly available RNA-seq data from The Cancer Genome Atlas Liver Hepatocellular Carcinoma (TCGA-LIHC) cohort were used to examine correlations between *SUV39H2* and AHR-responsive gene expression (Cancer Genome Atlas Research Network 2017).

Because MeRIP-seq and matched input RNA-seq were performed without biological replication, these analyses were considered exploratory and used primarily to identify candidate TCDD-responsive transcripts and m⁶A-enriched regions. Sequencing-derived statistical estimates were not interpreted as confirmatory evidence. Selected targets, including *CYP1A1*, *CYP1A2*, *AHRR*, *SUV39H2*, and *SMN1*, were subsequently examined in independent biological experiments using MeRIP-qPCR, RT-qPCR, immunoblotting, and mRNA stability assays. Additional SUV39H2 gain- and loss-of-function studies were performed to investigate its role in AHR signaling. Mechanistic conclusions were based on these independent validation and functional experiments rather than sequencing results alone.

### Methylated RNA immunoprecipitation followed by quantitative PCR (MeRIP-qPCR)

To validate TCDD-induced changes in N⁶-methyladenosine (m⁶A) enrichment identified by MeRIP-seq, MeRIP-qPCR was performed using the EpiQuik CUT&RUN m⁶A RNA Enrichment (MeRIP) Kit (EpigenTek, Cat. No. P-9018, Farmingdale, NY, USA) according to the manufacturer’s instructions. Total RNA was isolated from HepG2 cells treated with vehicle (DMSO) or 10 nM TCDD for 5 h. RNA samples were incubated with an anti-m⁶A antibody and affinity beads in immunocapture buffer for 90 min at room temperature. Following immunocapture, RNA cleavage was performed using the supplied nuclear digestion enhancer and cleavage enzyme mix to remove unbound RNA regions while preserving antibody-associated m⁶A-enriched fragments. After washing, captured RNA fragments were released by proteinase K digestion at 55 °C for 15 min and purified using magnetic RNA-binding beads. Purified m⁶A-enriched RNA and corresponding input RNA were reverse-transcribed into complementary DNA and analyzed by quantitative PCR using gene-specific primers. Relative m⁶A enrichment was determined by normalizing immunoprecipitated RNA signals to the corresponding input RNA signals and comparing TCDD-treated samples with vehicle controls. Selected transcripts included *CYP1A1*, *CYP1A2*, *AHRR*, *SUV39H2*, and *SMN1*. Primer sequences are provided in **Supplementary Table S3**.

### Small interfering RNA transfection and SUV39H2 overexpression

For loss-of-function experiments, cells were transfected with small interfering RNA targeting SUV39H2 (siSUV39H2; SASI_Hs01_00101226 and SASI_Hs02_00357085, Sigma-Aldrich, St. Louis, MO, USA). A non-targeting siRNA (Universal Negative Control, Cat. No. SIC001; Sigma-Aldrich) served as the negative control. Cells were incubated for 48 h following transfection before subsequent treatments or analyses. For gain-of-function experiments, cells were transfected with an SUV39H2 expression plasmid obtained from Addgene (Plasmid #89135) using Lipofectamine 3000 (Thermo Fisher Scientific, Waltham, MA, USA). Plasmid DNA was mixed with P3000 reagent and Lipofectamine 3000 in Opti-MEM according to the manufacturer’s instructions. Cells transfected with the corresponding empty vector served as controls. Following 24 h of transfection, cells were subjected to the indicated treatments or harvested for subsequent analyses. Changes in SUV39H2 expression were evaluated using RT-qPCR and/or immunoblotting.

### mRNA stability assay

To evaluate the effects of TCDD on mRNA stability, cells were treated with vehicle or TCDD under the indicated experimental conditions, followed by transcriptional inhibition using actinomycin D (ActD; 5 µg/mL) or flavopiridol (500 nM). Total RNA was isolated at 0, 4, and 8 h following transcriptional inhibition using RNAzol RT (Sigma-Aldrich). Transcript abundance was determined by quantitative reverse transcription PCR (RT-qPCR) and expressed relative to the corresponding 0-h value. mRNA decay curves were generated using GraphPad Prism, and apparent mRNA half-lives were estimated by fitting the data to a one-phase exponential decay model (Bensaude 2011; Lai et al. 2019). These experiments were used to assess the stability of selected AHR-responsive and m⁶A-enriched transcripts following TCDD exposure.

### Quantitative reverse transcription PCR (RT-qPCR)

Total RNA was isolated from HepG2 and Hep3B cells or liver tissue using RNAzol RT (Sigma-Aldrich) according to the manufacturer’s instructions. One microgram of total RNA was reverse-transcribed into complementary DNA (cDNA) using the High-Capacity cDNA Reverse Transcription Kit (Applied Biosystems, Thermo Fisher Scientific). Quantitative PCR was performed using SYBR Green Master Mix (Bio-Rad Laboratories, Hercules, CA, USA) on a QuantStudio 5 Real-Time PCR System (Thermo Fisher Scientific). Relative gene expression was calculated using the 2−ΔΔCt method, with β-actin (*ACTB*) as the internal reference gene. Primer sequences are provided in **Supplementary Table S3**. RT-qPCR experiments were independently repeated three times.

### Immunoblotting

Total protein lysates were prepared using radioimmunoprecipitation assay (RIPA) buffer (Thermo Fisher Scientific). Protein concentrations were determined using a bicinchoninic acid (BCA) assay. Equal amounts of protein were separated by sodium dodecyl sulfate–polyacrylamide gel electrophoresis (SDS–PAGE) and transferred onto polyvinylidene difluoride (PVDF) membranes (Millipore). Membranes were blocked using Proteintech Blotting Accelerator or nonfat dry milk and incubated with the indicated primary antibodies overnight at 4 °C. After washing, membranes were incubated with horseradish peroxidase (HRP)-conjugated secondary antibodies. Protein bands were detected using enhanced chemiluminescence (ECL) and imaged using an Odyssey Imaging System (LI-COR Biosciences). β-Actin was used as a loading control. Primary antibodies against β-actin, AHRR, CYP1A1, CYP1A2, CYP1B1, GCNT1, TTBK1, RAPSN, PLIN5, ROR1, CCR7, SUV39H2, and SMN were obtained from Cell Signaling Technology Aviva Systems Biology, Epigentek, or Proteintech. Antibody sources, and catalog numbers are provided in **Supplementary Table S4.**

### Luciferase reporter assay

AHR-dependent transcriptional activity was evaluated using a dioxin-responsive element (DRE)-driven firefly luciferase reporter assay. Cells were transfected with the DRE-luciferase reporter together with an SUV39H2 expression plasmid or siRNA targeting SUV39H2, as indicated. Following transfection, cells were treated with TCDD, an SUV39H2 inhibitor, or the corresponding vehicle controls under the specified experimental conditions. To further investigate whether SUV39H2 exhibits TCDD-responsive transcriptional regulatory activity, a GAL4-based luciferase reporter assay was performed. Cells were co-transfected with a GAL4-responsive luciferase reporter and a plasmid encoding a GAL4 DNA-binding domain–SUV39H2 fusion protein (GAL4-SUV39H2-L; Addgene plasmid #89136). Following transfection, cells were treated with TCDD or vehicle to assess the effects of AHR activation on GAL4-SUV39H2-mediated reporter activity. The corresponding control plasmids were included as indicated. For luciferase measurements, cells were lysed using Passive Lysis Buffer (Promega, Madison, WI, USA), and lysates were stored at −80 °C until analysis. Firefly and Renilla luciferase activities were measured using the Dual-Glo Luciferase Assay System (Promega) according to the manufacturer’s instructions. Because Renilla luciferase activity varied substantially across experimental conditions, firefly luciferase activity was normalized to total protein concentration, as determined using a bicinchoninic acid (BCA) assay, rather than to Renilla luciferase activity. Relative luciferase activity was expressed as protein-normalized firefly luminescence relative to the corresponding control group. Data were analyzed using GraphPad Prism.

### Cleavage under targets and release using nuclease followed by quantitative PCR (CUT&RUN-qPCR)

CUT&RUN assays were performed using the CUT&RUN Kit (Cell Signaling Technology, Danvers, MA, USA) according to the manufacturer’s instructions, with modifications. Cells were crosslinked with 0.1% formaldehyde for 4 min at room temperature, followed by quenching with glycine. Equal amounts of cells (∼200,000) were incubated overnight at 4 °C with antibodies against AHR or SUV39H2, or normal IgG as a negative control, in the presence of affinity beads. Following antibody incubation, samples were treated with nuclear digestion enhancer and cleavage enzyme mix. After washing, released DNA fragments were recovered by proteinase K treatment at 60 °C for 15 min and purified using magnetic DNA-binding beads. Because of the low yield of recovered DNA, sequencing libraries were prepared and amplified using Multiplex Oligos for Illumina Systems (Cell Signaling Technology). Amplified libraries were subjected to 0.9× size selection using AMPure XP beads (Beckman Coulter). The resulting DNA libraries were analyzed by quantitative PCR (qPCR) using primers targeting the *CYP1A1* enhancer region. CUT&RUN enrichment was calculated relative to the spike-in DNA reference used in the assay. Antibodies against AHR and SUV39H2 are listed in **Supplementary Table S4**, and the corresponding qPCR primers are provided in **Supplementary Table S3.**

### Animal experiments

C57BL/6J male mice (8- 12 weeks old) were used to evaluate the effects of TCDD exposure and pharmacological inhibition of SUV39H2 *in vivo*. Mice were randomly assigned to four treatment groups (n = 3–5 per group): vehicle control, TCDD alone, SUV39H2 inhibitor alone, and TCDD plus SUV39H2 inhibitor. Mice were pretreated with the SUV39H2 inhibitor OTS186935 (10 mg/kg, intraperitoneally i.p.) before administration of TCDD (10 µg/kg, i.p.). Around sixteen hours after TCDD administration, liver tissues were collected for RNA and protein extraction, followed by RT-qPCR and immunoblot analyses. Mice were housed in a pathogen-free facility under a 12-h light/dark cycle with standard chow and water provided ad libitum. All animal procedures were conducted in accordance with protocols approved by the Institutional Animal Care and Use Committee (IACUC) at the University of Arkansas for Medical Sciences.

### Statistical analysis

Statistical analyses of independently replicated experiments were performed using GraphPad Prism 10 (GraphPad Software, Boston, MA, USA). Data are presented as mean ± standard deviation (SD). Unless otherwise indicated, experimental validation assays were independently repeated at least three times. Differences among three or more groups were evaluated using one-way analysis of variance (ANOVA), whereas two-way ANOVA was applied to experiments involving two independent factors, as appropriate. The specific statistical tests and numbers of biological replicates are indicated in the corresponding figure legends. Statistical significance is indicated as *P* < 0.05 (\**), P < 0.01 (**), P < 0.001 (*\*\*\*), and *P* < 0.0001 (****). The MeRIP-seq discovery experiment included one biological sample per condition and was analyzed using exploratory statistical approaches. Sequencing-derived P values were used for candidate prioritization rather than confirmatory inference.

## Data availability

The MeRIP-seq datasets generated in this study have been deposited in the National Center for Biotechnology Information Gene Expression Omnibus (NCBI GEO) under accession number GSE351926 (BioProject PRJNA1549269). The deposited dataset includes raw sequencing data and processed gene expression data from vehicle- and TCDD-treated HepG2 cells.

## Results

### TCDD exposure alters the transcriptome-wide m⁶A landscape in HepG2 cells

To characterize the effects of TCDD exposure on RNA N⁶-methyladenosine (m⁶A) modification, we performed methylated RNA immunoprecipitation followed by sequencing (MeRIP-seq) in HepG2 cells treated with vehicle or 10 nM TCDD for 5 h. Transcriptome-wide analysis identified 14,265 genes associated with m⁶A-enriched regions in control cells and 14,177 genes in TCDD-treated cells. Among these, 12,594 genes were shared between the two conditions, whereas 1,671 and 1,583 genes were uniquely identified in control and TCDD-treated cells, respectively (**Fig. 1A**).

**Figure 1.**
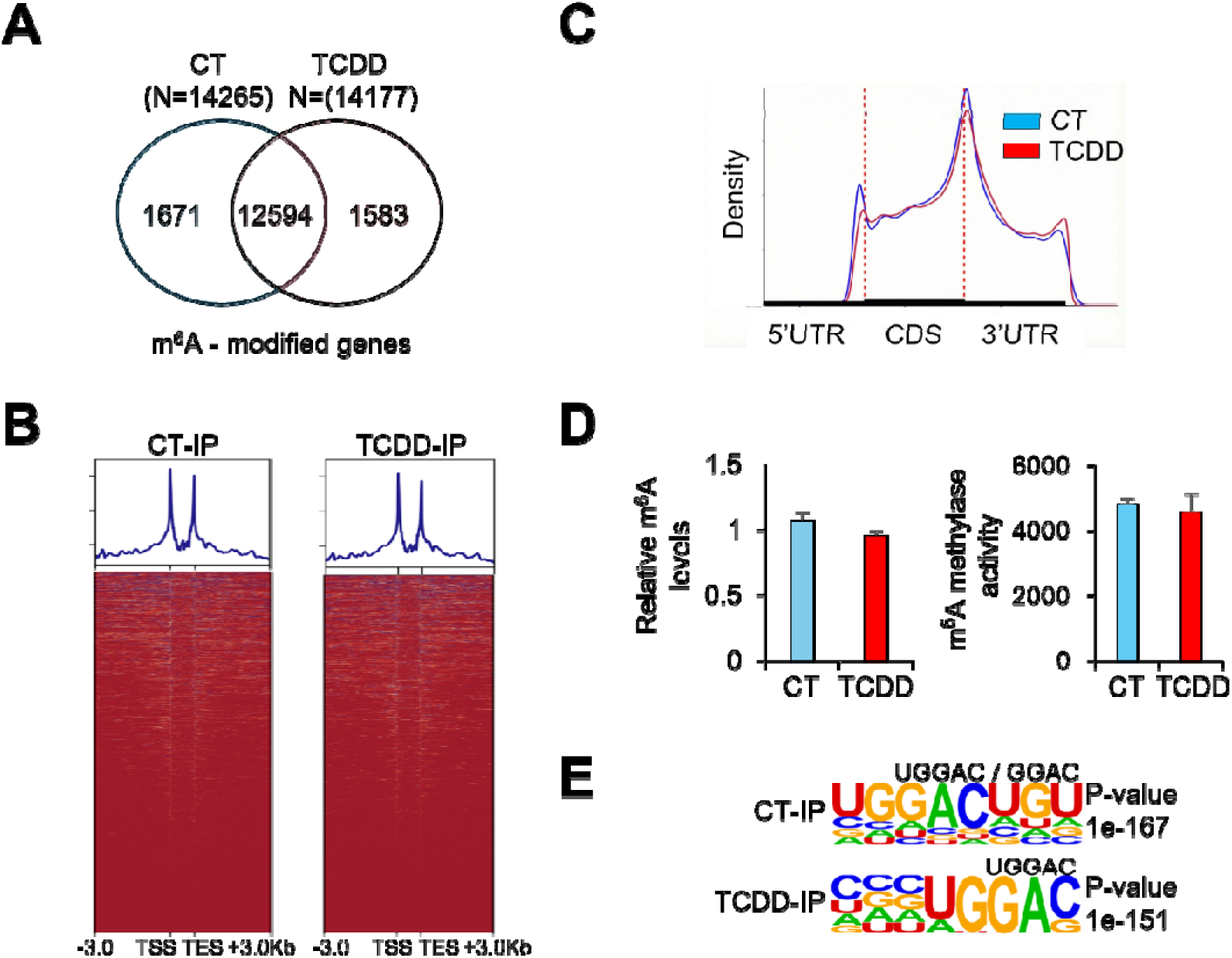
Transcriptome-wide profiling of m⁶A RNA modifications in control and TCDD-treated HepG2 cells. HepG2 cells were treated with vehicle (CT; DMSO) or 10 nM TCDD for 5 h, followed by methylated RNA immunoprecipitation sequencing (MeRIP-seq) to characterize transcriptome-wide m⁶A enrichment. **(A)** Venn diagram showing the numbers of m⁶A-enriched peaks identified in control and TCDD-treated cells. A total of 14,265 and 14,177 peaks were detected in the respective groups, including 12,594 shared peaks and 1,671 and 1,583 condition-specific peaks, respectively. **(B)** Metagene profiles and corresponding heatmaps showing the distribution of m⁶A-enriched sequencing signals across annotated gene regions, including the transcription start site (TSS), transcription end site (TES), and 3-kb upstream and downstream flanking regions. **(C)** Metagene density profiles illustrating the relative distribution of m⁶A enrichment across the 5′ untranslated region (5′UTR), coding sequence (CDS), and 3′ untranslated region (3′UTR) in control and TCDD-treated cells. Dashed lines indicate transcript-region boundaries. **(D)** Relative global m⁶A levels and m⁶A methyltransferase activity in control and TCDD-treated cells. Data are presented as mean ± SD from three independent experiments. **(E)** Sequence motif analysis of m⁶A-enriched regions in control and TCDD-treated cells, showing enrichment of canonical GGAC-containing motifs with corresponding motif-enrichment *P* values. Additional experimental and bioinformatics details are provided in Materials and Methods.

Metagene profiling and heatmap analysis revealed broadly similar distributions of m⁶A-enriched regions across the transcriptome in both groups (**Fig. 1B**). Analysis of m⁶A peak distribution along transcript regions showed preferential enrichment near the coding sequence (CDS)–3′ untranslated region (3′UTR) boundary, consistent with the characteristic distribution of m⁶A modifications (**Fig. 1C**). Despite differences in the genes associated with detectable m⁶A peaks, global m⁶A levels and m⁶A methyltransferase activity remained largely unchanged following TCDD exposure (**Fig. 1D**).

Sequence motif analysis further revealed strong enrichment of canonical m⁶A-associated motifs containing the GGAC core sequence in both vehicle- and TCDD-treated groups (**Fig. 1E**). These motifs are consistent with the conserved DRACH consensus sequence (D = A, G, or U; R = A or G; H = A, C, or U), a preferential sequence context for m⁶A deposition (Dominissini et al. 2012; Meyer et al. 2012). The preservation of these motifs suggests that TCDD exposure does not substantially alter the characteristic sequence context of m⁶A modification.

Collectively, these findings demonstrate that TCDD exposure is associated with changes in the transcriptome-wide landscape of detectable m⁶A-enriched regions without substantially altering global m⁶A abundance, methyltransferase activity, or the overall positional distribution of m⁶A modifications. These observations suggest that TCDD may preferentially influence m⁶A enrichment at selected transcripts rather than inducing widespread changes in global RNA methylation.

### TCDD induces distinct changes in m⁶A enrichment and gene expression in HepG2 cells

To further characterize TCDD-induced changes in RNA methylation and their relationship to gene expression, we performed differential m⁶A peak and transcriptome analyses in HepG2 cells following exposure to 10 nM TCDD for 5 h. Differential peak analysis identified both increased and decreased m⁶A enrichment following TCDD treatment (**Fig. 2A**). Notably, increased m⁶A enrichment was observed at regions associated with *SUV39H2*, *SMN1*, and *ZNF322*, whereas decreased enrichment was detected at regions associated with *EIF3C*, *LSM12*, and *SNRPG*. Peaks associated with *SUV39H2* and *SMN1* were among the top 10 regions showing increased m⁶A enrichment, with peaks annotated to the 5′ UTR of *SUV39H2* and the 3′ UTR of *SMN1* (**Supplementary Table S1**). These regions are also illustrated in the IGV tracks (**Supplementary Fig. S2**). These findings indicate that TCDD exposure induces transcript-associated changes in m⁶A enrichment despite the absence of substantial alterations in global m⁶A levels.

**Figure 2.**
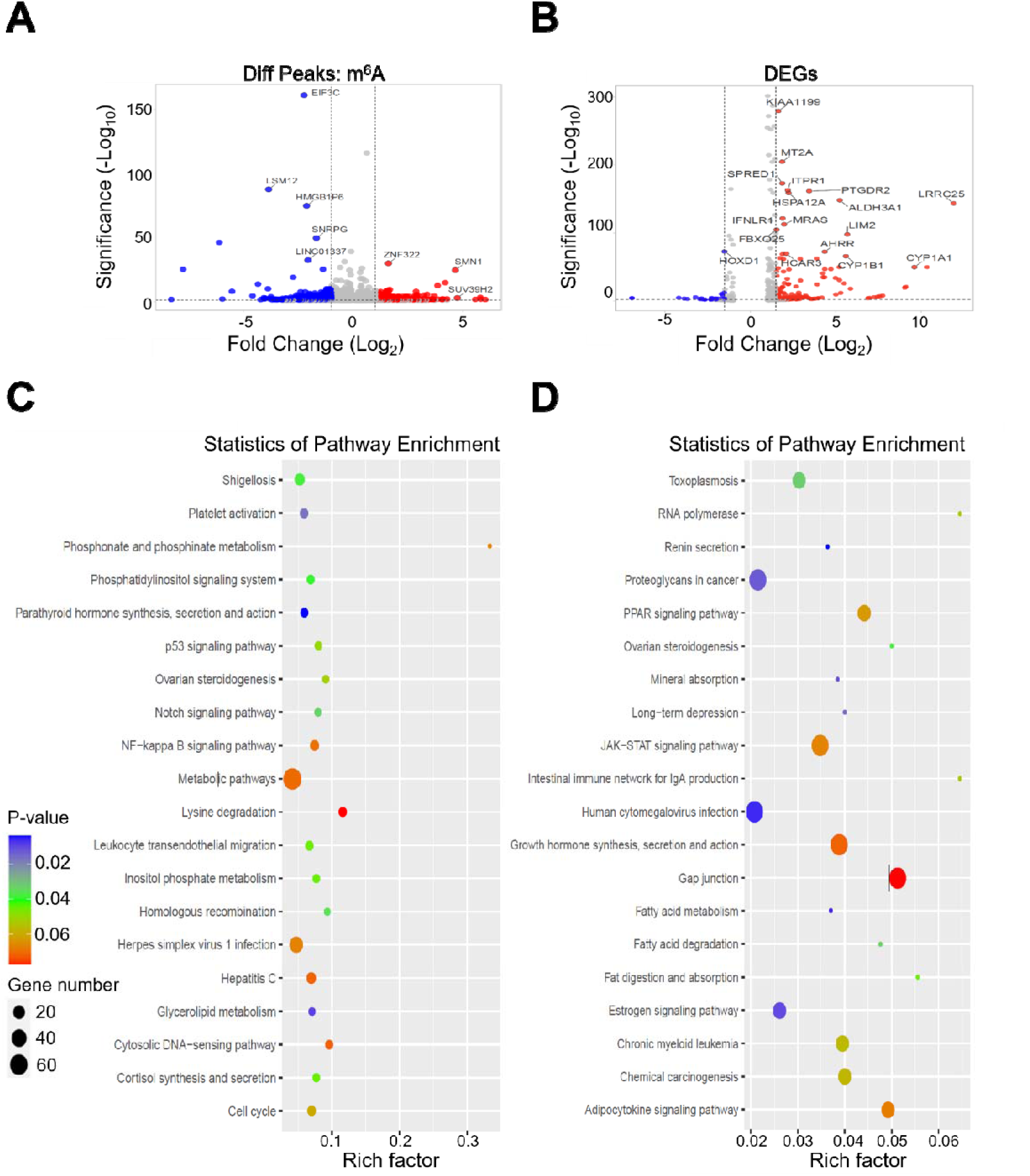
Transcriptome-wide changes in m⁶A enrichment and gene expression following TCDD exposure. HepG2 cells were treated with vehicle (DMSO) or 10 nM TCDD for 5 h, followed by MeRIP-seq and matched input RNA-seq analyses. **(A)** Volcano plot showing differential m⁶A peak enrichment between vehicle- and TCDD-treated cells. Red and blue dots indicate peaks with increased and decreased m⁶A enrichment, respectively, while gray dots represent peaks not meeting the indicated statistical thresholds. Selected genes associated with differential m⁶A peaks are labeled. **(B)** Volcano plot showing changes in gene expression between vehicle- and TCDD-treated cells. Red and blue dots indicate upregulated and downregulated genes, respectively, while gray dots represent genes not meeting the indicated statistical thresholds. Selected TCDD-responsive genes, including *CYP1A1*, *CYP1B1*, and *AHRR*, are labeled. **(C)** Kyoto Encyclopedia of Genes and Genomes (KEGG) pathway enrichment analysis of genes associated with differential m⁶A peaks. **(D)** KEGG pathway enrichment analysis of candidate differentially expressed genes following TCDD exposure. In (C) and (D), dot size represents the number of genes associated with each pathway, dot color indicates the nominal enrichment *P* value, and the x-axis represents the rich factor (the proportion of genes associated with a given pathway relative to the total number of genes annotated to that pathway). Sequencing-based analyses were performed without biological replicates and are considered exploratory.

In parallel, differential gene expression analysis revealed pronounced transcriptional responses to TCDD, including marked induction of canonical AHR-responsive genes such as *CYP1A1*, *CYP1B1*, and *AHRR* (**Fig. 2B**), consistent with activation of AHR-dependent transcriptional signaling. The top 10 upregulated genes, ranked by log₂ fold change, are summarized (**Supplementary Table S2**).

To explore the biological processes associated with these molecular changes, we performed Kyoto Encyclopedia of Genes and Genomes (KEGG) pathway enrichment analyses. Genes associated with differentially enriched m⁶A peaks were represented in pathways related to metabolism, phosphatidylinositol signaling, p53 signaling, NF-κB signaling, and lysine degradation (**Fig. 2C**). In contrast, differentially expressed genes were associated with pathways involving PPAR signaling, fatty acid metabolism, JAK–STAT signaling, chemical carcinogenesis, and other cellular signaling processes (**Fig. 2D**).

The differences in pathway representation between genes associated with differential m⁶A enrichment and differentially expressed genes suggest that TCDD-induced changes in RNA methylation and transcript abundance may involve partially distinct regulatory processes. Together, these findings identify transcript-associated m⁶A remodeling as an additional molecular response to TCDD exposure and provide a foundation for investigating whether m⁶A modifications contribute to the post-transcriptional regulation of AHR-responsive genes.

### TCDD enhances m⁶A enrichment and prolongs the mRNA stability of canonical AHR-responsive genes

To investigate whether TCDD-induced m⁶A remodeling is associated with post-transcriptional regulation of AHR-responsive genes, we examined m⁶A enrichment and mRNA stability of three canonical AHR target genes, *CYP1A1*, *AHRR*, and *CYP1A2*, in HepG2 cells. Genome browser visualization of MeRIP-seq data revealed prominent m⁶A-enriched regions associated with *CYP1A1*, *AHRR*, and *CYP1A2* transcripts following TCDD exposure, whereas minimal enrichment signals were detected in vehicle-treated cells (**Fig. 3A**). Consistent with these observations, MeRIP-qPCR analysis demonstrated increased m⁶A enrichment of all three transcripts following TCDD treatment, with substantially greater enrichment at 24 h than at 6 h (**Fig. 3B**).

**Figure 3.**
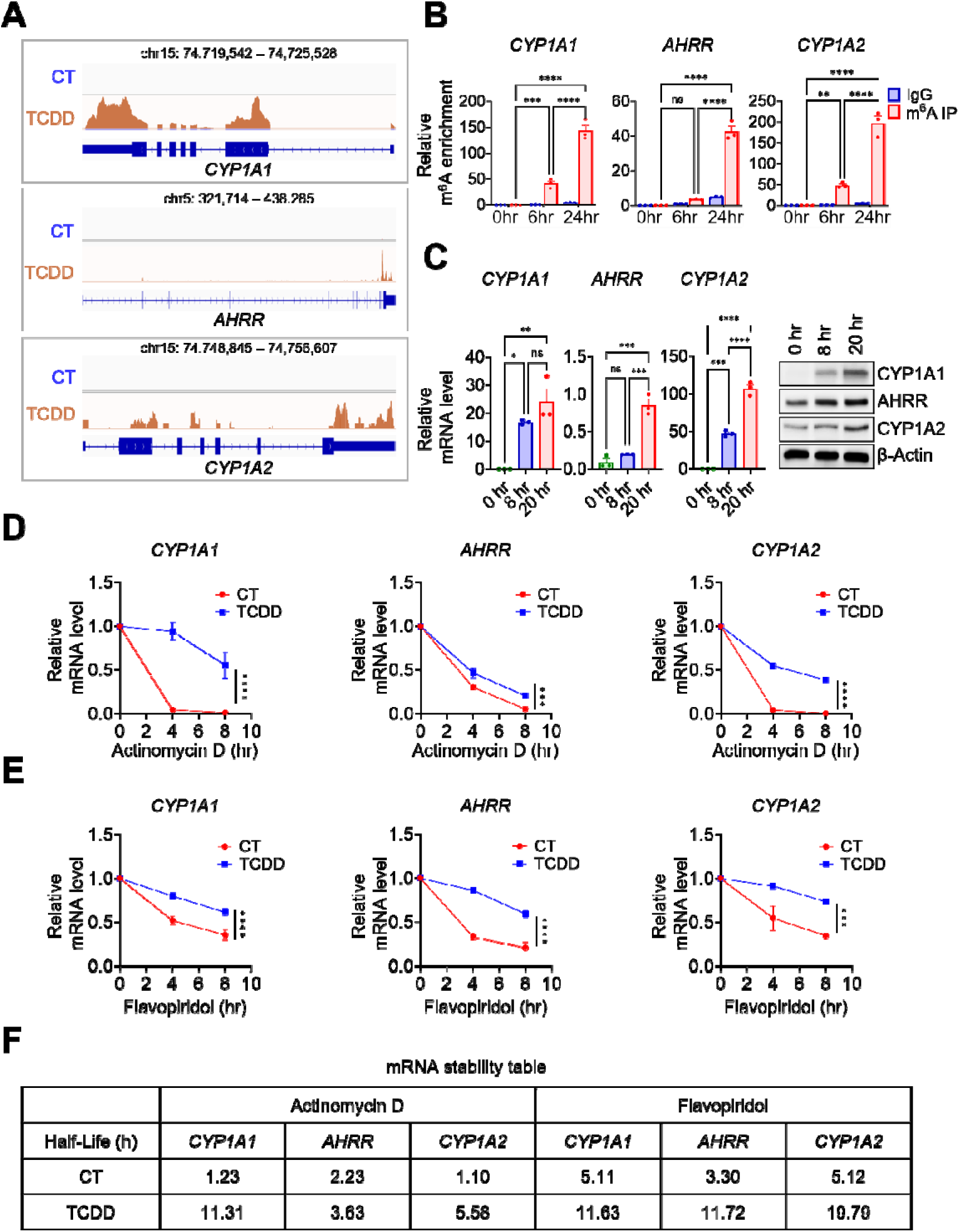
TCDD increases m⁶A enrichment and prolongs the mRNA stability of canonical AHR-responsive genes. **(A)** Integrative Genomics Viewer (IGV) tracks showing m⁶A-enriched regions associated with *CYP1A1*, *AHRR*, and *CYP1A2* transcripts in vehicle-treated control (CT) and TCDD-treated HepG2 cells. Genomic coordinates and annotated gene structures are indicated. **(B)** MeRIP-qPCR analysis of m⁶A enrichment in *CYP1A1*, *AHRR*, and *CYP1A2* transcripts following TCDD exposure for 0, 6, and 24 h. Enrichment was normalized to the corresponding input RNA, with IgG immunoprecipitation serving as a negative control. **(C)** RT-qPCR analysis of *CYP1A1*, *AHRR*, and *CYP1A2* mRNA expression following TCDD exposure for 0, 8, and 20 h. Transcript levels were normalized to *ACTB* expression. Representative immunoblots show the corresponding protein abundance, with β-actin serving as a loading control. **(D, E)** mRNA decay profiles of *CYP1A1*, *AHRR*, and *CYP1A2* in vehicle- and TCDD-treated HepG2 cells following transcriptional inhibition with actinomycin D (D) or flavopiridol (E). Transcript abundance was determined by RT-qPCR at 0, 4, and 8 h after transcriptional inhibition and normalized to the corresponding 0-h value. **(F)** Summary of apparent mRNA half-lives (t₁/₂) for *CYP1A1*, *AHRR*, and *CYP1A2* following transcriptional inhibition with actinomycin D or flavopiridol. Half-lives were estimated by fitting mRNA decay curves to a one-phase exponential decay model and are expressed in hours. Quantitative data are presented as mean ± SD from three independent experiments. Statistical significance is indicated as *P* < 0.05 (*), *P* < 0.01 (**), *P* < 0.001 (***), and *P* < 0.0001 (****); ns, not significant. Statistical tests are described in Materials and Methods.

We next examined the temporal expression patterns of these AHR-responsive genes. RT-qPCR analysis demonstrated that TCDD increased *CYP1A1*, *AHRR*, and *CYP1A2* mRNA expression at 8 and 20 h, with transcript-specific differences in the magnitude and kinetics of induction (**Fig. 3C**). Immunoblot analysis further demonstrated increased CYP1A1, AHRR, and CYP1A2 protein abundance following TCDD exposure (**Fig. 3C**). Together, these findings indicate that increased m⁶A enrichment of canonical AHR-responsive transcripts accompanies their induction at both the mRNA and protein levels.

To determine whether TCDD exposure also affects the stability of these transcripts, we examined mRNA decay following transcriptional inhibition using actinomycin D and flavopiridol, which inhibit transcription through distinct mechanisms (Bensaude 2011). Following actinomycin D treatment, *CYP1A1*, *AHRR*, and *CYP1A2* transcripts exhibited more rapid decay in vehicle-treated cells than in TCDD-treated cells (**Fig. 3D**). The estimated mRNA half-lives increased from 1.23 to 11.31 h for *CYP1A1*, 2.23 to 3.63 h for *AHRR*, and 1.10 to 5.58 h for *CYP1A2* following TCDD exposure (**Fig. 3F**). Consistent with these findings, flavopiridol-mediated transcriptional inhibition also revealed slower transcript decay in TCDD-treated cells (**Fig. 3E**). The estimated half-lives increased from 5.11 to 11.63 h for *CYP1A1*, 3.30 to 11.72 h for *AHRR*, and 5.12 to 19.79 h for *CYP1A2* (**Fig. 3F**). Although the absolute half-life estimates differed between the two transcriptional inhibitors, both approaches consistently indicated increased transcript stability following TCDD exposure.

Collectively, these findings demonstrate that TCDD exposure increases m⁶A enrichment of canonical AHR-responsive transcripts and is associated with prolonged mRNA stability. Previous studies have established that m⁶A modifications can influence mRNA turnover through distinct reader-dependent mechanisms, including transcript destabilization and stabilization (Wang et al. 2014; Huang et al. 2018). These observations suggest that post-transcriptional mechanisms may contribute to the sustained expression of AHR target genes following TCDD exposure, although the direct contribution of m⁶A modification to transcript stabilization remains to be established.

### Integrated transcriptomic analysis identifies distinct relationships between m⁶A enrichment and gene expression following TCDD exposure

To investigate the relationship between TCDD-induced m⁶A remodeling and gene expression, we integrated MeRIP-seq and RNA-seq datasets from TCDD-treated HepG2 cells. This analysis identified four groups of genes exhibiting distinct patterns of m⁶A enrichment and transcript abundance (**Fig. 4A**). Applying thresholds of greater than 1.25-fold change in m⁶A enrichment and greater than 2-fold change in mRNA expression, 146 genes exhibited concomitant increases in m⁶A enrichment and transcript abundance, including the canonical AHR-responsive genes *CYP1A1*, *CYP1A2*, and *AHRR*. In contrast, 79 genes exhibited decreased m⁶A enrichment despite increased mRNA expression, including *ROR1* and *CCR7*. Among genes with reduced mRNA expression, 41 exhibited decreased m⁶A enrichment, including *GCNT1* and *TTBK1*, whereas 60 exhibited increased m⁶A enrichment, including *PLIN5* and *RAPSN*. Genome browser tracks for the six selected genes are presented (**Supplementary Fig. S1**).

**Figure 4.**
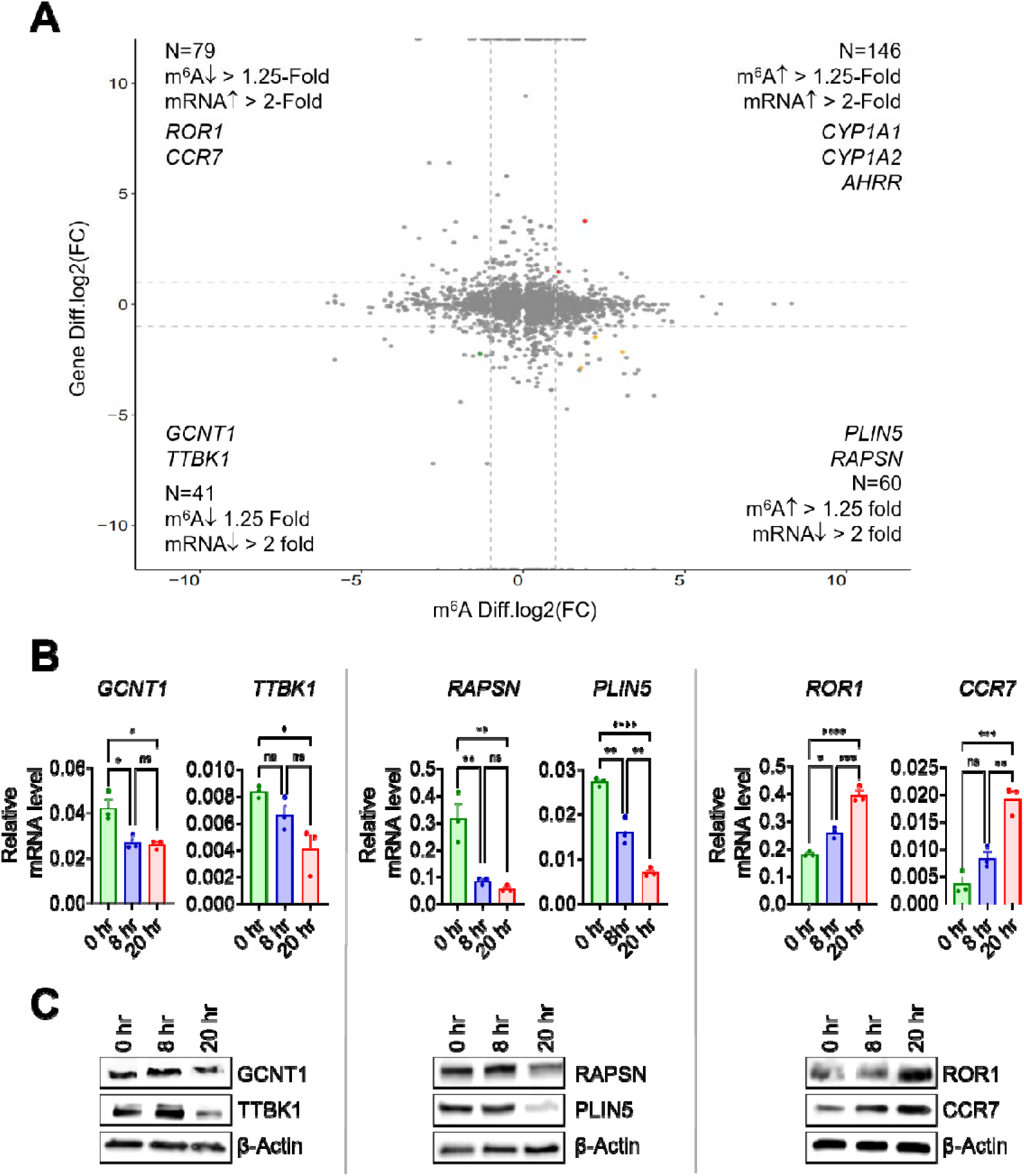
Integrated analysis reveals distinct patterns of m⁶A enrichment and gene expression following TCDD exposure. **(A)** Four-quadrant scatter plot showing the relationship between changes in m⁶A enrichment and mRNA expression in TCDD-treated HepG2 cells relative to vehicle-treated controls. The x-axis represents the log₂ fold change in m⁶A enrichment, and the y-axis represents the log₂ fold change in mRNA expression. Genes were classified into four groups using fold-change thresholds of >1.25 for m⁶A enrichment and >2 for mRNA expression: increased m⁶A/increased mRNA (n = 146), decreased m⁶A/increased mRNA (n = 79), decreased m⁶A/decreased mRNA (n = 41), and increased m⁶A/decreased mRNA (n = 60). Representative genes from each group are indicated. **(B)** RT-qPCR analysis of *GCNT1*, *TTBK1*, *RAPSN*, *PLIN5*, *ROR1*, and *CCR7* mRNA expression following TCDD exposure for 0, 8, and 20 h. Transcript levels were normalized to *ACTB* expression. **(C)** Representative immunoblots showing GCNT1, TTBK1, RAPSN, PLIN5, ROR1, and CCR7 protein abundance following TCDD exposure for 0, 8, and 20 h, with β-actin serving as a loading control. RT-qPCR data are presented as mean ± SD from three independent experiments. Statistical significance is indicated as *P* < 0.05 (*), *P* < 0.01 (**), *P* < 0.001 (***), and *P* < 0.0001 (****); ns, not significant. Statistical tests are described in Materials and Methods.

To examine the expression patterns of representative genes identified by the integrated analysis, we evaluated their mRNA and protein abundance following TCDD exposure. RT-qPCR analysis demonstrated significantly decreased *GCNT1*, *RAPSN*, and *PLIN5* mRNA expression, whereas *ROR1* and *CCR7* expression increased following TCDD treatment (**Fig. 4B**). *TTBK1* expression was also significantly reduced at 20 h. Immunoblot analysis further revealed time-dependent changes in the corresponding protein levels (**Fig. 4C**). In particular, GCNT1, TTBK1, RAPSN, and PLIN5 protein abundance decreased at 20 h, whereas ROR1 and CCR7 protein abundance increased.

Collectively, these findings demonstrate that TCDD exposure is associated with heterogeneous relationships between m⁶A enrichment and transcript abundance. Increased m⁶A enrichment was observed in association with both increased and decreased mRNA expression, indicating that m⁶A remodeling does not uniformly correlate with transcript upregulation. These observations suggest that the relationship between TCDD-associated m⁶A remodeling and gene expression is transcript-dependent and may reflect distinct post-transcriptional regulatory contexts.

### TCDD increases m⁶A enrichment and mRNA stability of SUV39H2 and SMN1 without altering steady-state transcript abundance

To further investigate the relationship between TCDD-induced m⁶A remodeling and post-transcriptional gene regulation, we examined *SUV39H2* and *SMN1*, two transcripts identified as exhibiting increased m⁶A enrichment in the transcriptome-wide analysis (**Fig. 2A**, **Supplementary Table S1**). RT-qPCR analysis revealed no significant changes in *SUV39H2* or *SMN1* mRNA abundance following TCDD exposure for 8 or 20 h (**Fig. 5A**). In contrast, MeRIP-qPCR analysis demonstrated marked, time-dependent increases in m⁶A enrichment of both transcripts following TCDD exposure for 6 and 24 h (**Fig. 5B**). These findings indicate that TCDD-associated increases in m⁶A enrichment can occur independently of detectable changes in steady-state transcript abundance. Despite the absence of significant changes in mRNA expression, immunoblot analysis revealed increased SUV39H2 and SMN1 protein abundance following TCDD exposure (**Fig. 5C**), suggesting that post-transcriptional mechanisms may contribute to their regulation.

**Figure 5.**
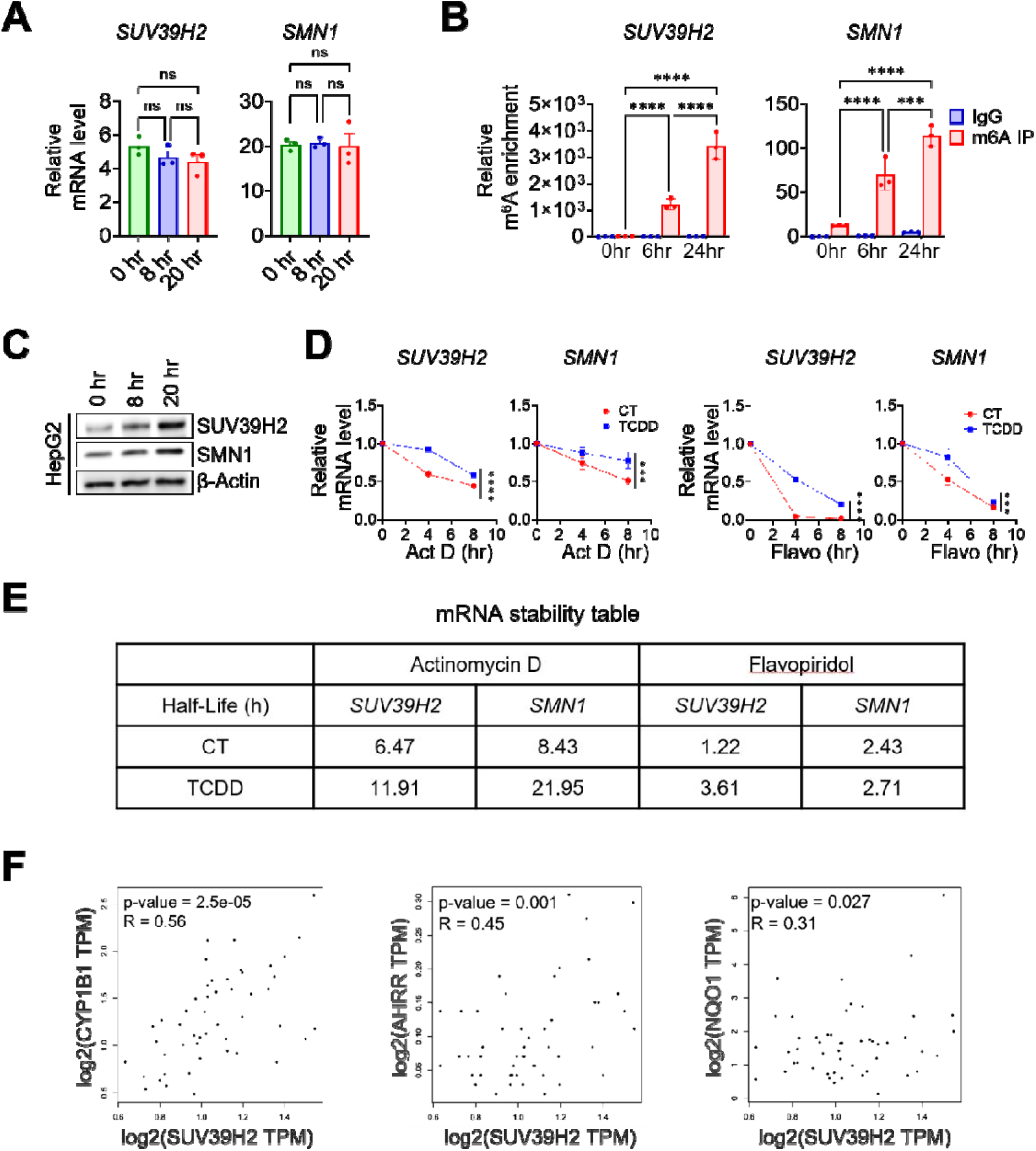
TCDD enhances m⁶A enrichment and alters the post-transcriptional regulation of SUV39H2 and SMN1 in HepG2 cells. **(A)** RT-qPCR analysis of *SUV39H2* and *SMN1* mRNA expression in HepG2 cells following TCDD exposure for 0, 8, and 20 h. Transcript levels were normalized to *ACTB* expression. **(B)** MeRIP-qPCR analysis of m⁶A enrichment in *SUV39H2* and *SMN1* transcripts following TCDD exposure for 0, 6, and 24 h. Enrichment was normalized to the corresponding input RNA, with IgG immunoprecipitation serving as a negative control. **(C)** Representative immunoblots showing SUV39H2 and SMN1 protein abundance following TCDD exposure for 0, 8, and 20 h, with β-actin serving as a loading control. **(D)** mRNA decay profiles of *SUV39H2* and *SMN1* in vehicle-treated control (CT) and TCDD-treated HepG2 cells following transcriptional inhibition with actinomycin D (left) or flavopiridol (right). Relative transcript abundance was determined by RT-qPCR at 0, 4, and 8 h after transcriptional inhibition and normalized to the corresponding 0-h value. **(E)** Summary of apparent mRNA half-lives (t₁/₂) for *SUV39H2* and *SMN1* following transcriptional inhibition with actinomycin D or flavopiridol. Half-lives were estimated by fitting mRNA decay curves to a one-phase exponential decay model and are expressed in hours. **(F)** Correlation analysis of *SUV39H2* expression with *CYP1B1*, *AHRR*, and *NQO1* expression using transcriptomic data from the TCGA liver hepatocellular carcinoma (TCGA-LIHC) cohort. Correlation coefficients (R) and corresponding *P* values are indicated. Quantitative experimental data are presented as mean ± SD from three independent biological replicates (n = 3). Statistical significance is indicated as *P* < 0.05 (*), *P* < 0.01 (**), *P* < 0.001 (***), and *P* < 0.0001 (****); ns, not significant. Statistical methods are described in Materials and Methods.

To determine whether TCDD exposure affects the stability of these transcripts, we examined mRNA decay following transcriptional inhibition with actinomycin D or flavopiridol. Both *SUV39H2* and *SMN1* exhibited slower decay in TCDD-treated cells than in controls following actinomycin D treatment (**Fig. 5D**). The estimated mRNA half-lives increased from 6.47 to 11.91 h for *SUV39H2* and from 8.43 to 21.95 h for *SMN1* (**Fig. 5E**). Consistent with these findings, flavopiridol-mediated transcriptional inhibition revealed prolonged *SUV39H2* mRNA stability, with an estimated half-life increase from 1.22 to 3.61 h. In contrast, *SMN1* exhibited only a modest increase in estimated half-life, from 2.43 to 2.71 h (**Fig. 5D, E**). Thus, increased apparent transcript stability following TCDD exposure was more consistently observed for *SUV39H2* than for *SMN1* across the two transcriptional inhibition approaches.

SUV39H2 is a histone H3K9 methyltransferase primarily associated with transcriptional repression but has also been implicated in context-dependent regulation of nuclear receptor activity (Askew et al. 2017). Given its potential involvement in transcriptional regulation, we next examined the relationship between *SUV39H2* expression and AHR-responsive genes using transcriptomic data from the TCGA liver hepatocellular carcinoma (TCGA-LIHC) cohort. Correlation analysis revealed significant positive associations between *SUV39H2* and *CYP1B1* (R = 0.56, *P* = 2.5 × 10⁻⁵), *AHRR* (R = 0.45, *P* = 0.001), and *NQO1* (R = 0.31, *P* = 0.027) (**Fig. 5F**). These associations suggest a potential relationship between SUV39H2 expression and AHR-responsive transcriptional programs, although they do not establish direct regulatory interactions.

Collectively, these findings demonstrate that TCDD exposure increases m⁶A enrichment of *SUV39H2* and *SMN1* transcripts without significantly altering their steady-state mRNA abundance. The accompanying increases in protein abundance and apparent mRNA stability are consistent with the involvement of post-transcriptional regulatory mechanisms. Previous studies have established that m⁶A modification can influence RNA stability, translation, and processing through distinct regulatory pathways (Wang et al. 2014; Wang et al. 2015; Roundtree et al. 2017; Huang et al. 2018). Accordingly, our findings raise the possibility that TCDD-associated m⁶A remodeling contributes to the post-transcriptional regulation of SUV39H2 and SMN1, although a direct causal relationship remains to be established. Together with the positive correlations between *SUV39H2* and AHR-responsive genes, these observations provided a rationale for further investigating the functional involvement of SUV39H2 in AHR signaling.

### SUV39H2 positively regulates AHR-dependent transcriptional activity

To investigate the functional involvement of SUV39H2 in AHR signaling, we examined the effects of SUV39H2 depletion and overexpression on AHR-dependent transcriptional responses in HepG2 and Hep3B cells. Transfection with two independent SUV39H2-targeting siRNAs reduced *SUV39H2* mRNA and protein expression in both cell lines (**Fig. 6A**). Conversely, ectopic expression of SUV39H2 markedly increased its mRNA and protein abundance (**Fig. 6B**), confirming the effectiveness of the loss- and gain-of-function approaches.

**Figure 6.**
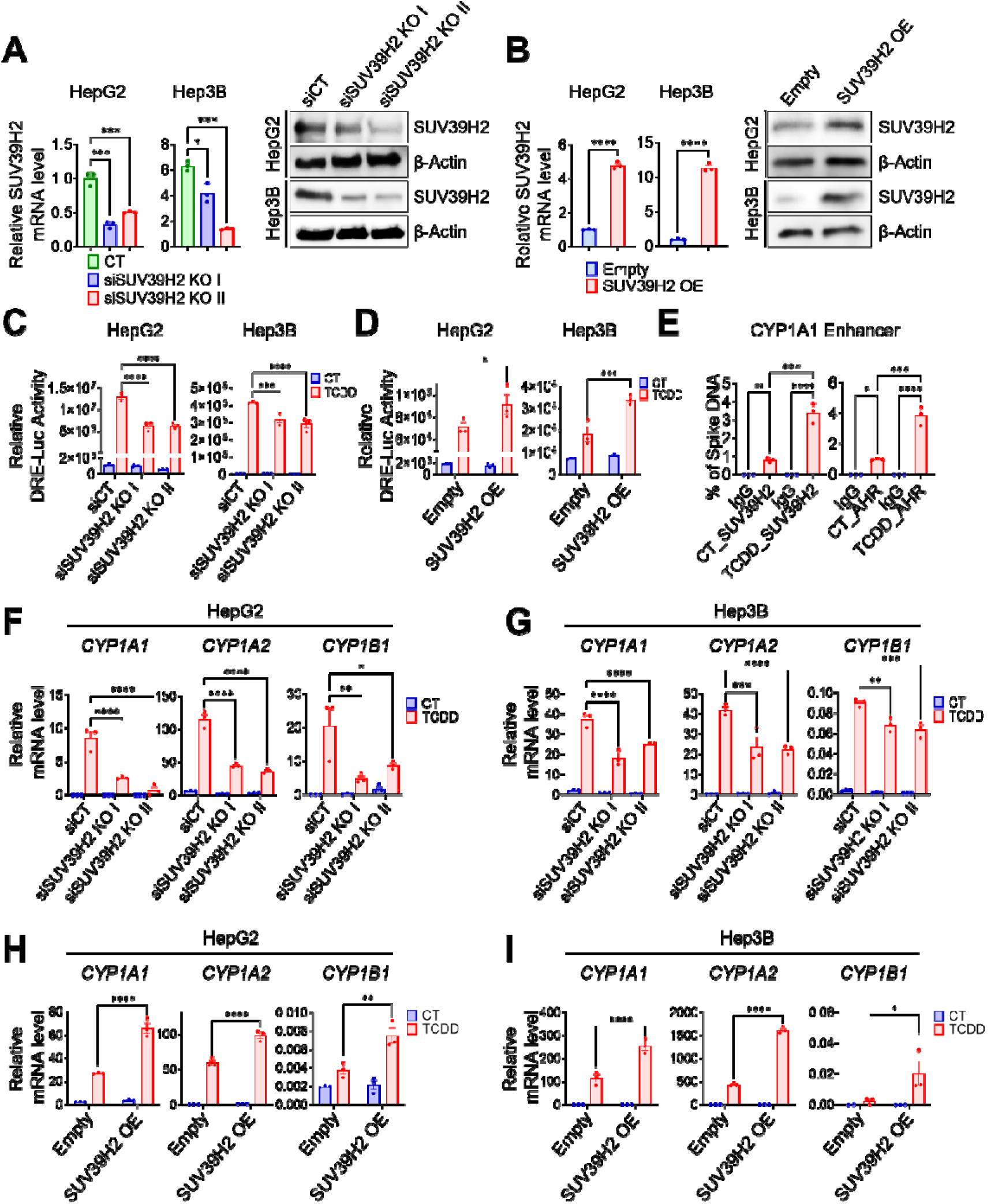
SUV39H2 enhances AHR-dependent transcriptional activity and target gene expression in hepatic cancer cells. **(A)** Validation of SUV39H2 knockdown in HepG2 and Hep3B cells transfected with control siRNA (siCT) or two independent SUV39H2-targeting siRNAs (siSUV39H2 KO I and KO II). *SUV39H2* mRNA and protein levels were assessed by RT-qPCR and immunoblotting, respectively, with β-actin serving as a protein loading control. **(B)** Validation of SUV39H2 overexpression in HepG2 and Hep3B cells transfected with empty vector or an SUV39H2 expression construct. *SUV39H2* mRNA and protein levels were assessed by RT-qPCR and immunoblotting, respectively, with β-actin serving as a loading control. **(C)** Dioxin response element (DRE)-driven luciferase reporter activity in HepG2 and Hep3B cells transfected with control or SUV39H2-targeting siRNAs and treated with vehicle (CT) or TCDD. **(D)** DRE-driven luciferase reporter activity in HepG2 and Hep3B cells following transfection with empty vector or an SUV39H2 expression construct and treatment with vehicle or TCDD. **(E)** CUT&RUN-qPCR analysis of SUV39H2 and AHR occupancy at the *CYP1A1* enhancer in vehicle-treated control (CT) and TCDD-treated HepG2 cells. Antibodies against SUV39H2 or AHR were used for CUT&RUN, with IgG serving as a negative control. Enrichment was expressed as the percentage of spike-in DNA. **(F, G)** RT-qPCR analysis of *CYP1A1*, *CYP1A2*, and *CYP1B1* mRNA expression following SUV39H2 knockdown in HepG2 (F) and Hep3B (G) cells treated with vehicle or TCDD. **(H, I)** RT-qPCR analysis of *CYP1A1*, *CYP1A2*, and *CYP1B1* mRNA expression following SUV39H2 overexpression in HepG2 (H) and Hep3B (I) cells treated with vehicle or TCDD. Quantitative data are presented as mean ± SD from three independent biological replicates (n = 3). Statistical significance is indicated as *P* < 0.05 (*), *P* < 0.01 (**), *P* < 0.001 (***), and *P* < 0.0001 (****); ns, not significant. Statistical methods are described in Materials and Methods.

We next assessed the contribution of SUV39H2 to AHR transcriptional activity using a dioxin response element (DRE)-driven luciferase reporter assay. TCDD markedly increased reporter activity in control cells, whereas SUV39H2 depletion significantly attenuated TCDD-induced reporter activation in both HepG2 and Hep3B cells (**Fig. 6C**). Conversely, SUV39H2 overexpression further enhanced TCDD-induced reporter activity compared with empty-vector controls in both cell lines (**Fig. 6D**). These complementary findings indicate that SUV39H2 positively modulates AHR-dependent reporter activity.

To determine whether SUV39H2 is recruited to AHR-responsive regulatory regions, we performed CUT&RUN followed by quantitative PCR (CUT&RUN-qPCR) targeting the *CYP1A1* enhancer, a well-characterized regulatory region involved in AHR-mediated transcription (Matthews et al. 2005). TCDD exposure significantly increased SUV39H2 occupancy at this region compared with vehicle-treated controls (**Fig. 6E**). Similarly, TCDD enhanced AHR occupancy at the same enhancer region. These findings demonstrate TCDD-induced enrichment of both SUV39H2 and AHR at the *CYP1A1* enhancer, supporting a potential role for SUV39H2 in AHR-mediated transcriptional regulation.

We further examined whether SUV39H2 influences endogenous AHR target gene expression. SUV39H2 depletion attenuated TCDD-induced *CYP1A1*, *CYP1A2*, and *CYP1B1* mRNA expression in HepG2 cells (**Fig. 6F**). Similar reductions were observed in Hep3B cells (**Fig. 6G**), although the magnitude of the response varied among individual genes and siRNAs. Conversely, SUV39H2 overexpression enhanced TCDD-induced expression of these AHR-responsive genes in both HepG2 and Hep3B cells (**Fig. 6H, I**). These findings demonstrate that SUV39H2 contributes to the magnitude of TCDD-induced AHR target gene expression in two hepatic cancer cell models.

Collectively, these results identify SUV39H2 as a positive modulator of TCDD-induced AHR transcriptional responses. The concordant effects of SUV39H2 depletion and overexpression on DRE-driven reporter activity and endogenous AHR target gene expression, together with increased SUV39H2 occupancy at the *CYP1A1* enhancer following TCDD exposure, support a potential coactivator-like role for SUV39H2 in AHR signaling. However, whether this activity requires SUV39H2-mediated histone methylation or direct interaction with AHR remains to be determined.

### Pharmacological inhibition of SUV39H2 attenuates TCDD-induced AHR signaling *in vitro* and *in vivo*

To further investigate the functional involvement of SUV39H2 in AHR signaling, we examined whether pharmacological inhibition of SUV39H2 affects TCDD-induced AHR transcriptional responses. In HepG2 cells, treatment with the SUV39H2 inhibitor at 500 nM or 1 µM attenuated TCDD-induced *CYP1A1*, *CYP1B1*, and *AHRR* mRNA expression. Similar inhibitory effects were observed in Hep3B cells at 1 and 3 µM, although the magnitude of inhibition varied among individual genes and concentrations (**Fig. 7A**). Consistent with these findings, SUV39H2 inhibition reduced TCDD-induced dioxin response element (DRE)-driven luciferase reporter activity in both HepG2 and Hep3B cells (**Fig. 7B**). In HepG2 cells, TCDD also increased GAL4-dependent reporter activity, whereas treatment with the SUV39H2 inhibitor markedly suppressed this response (**Fig. 7C**). Together, these findings indicate that pharmacological inhibition of SUV39H2 attenuates TCDD-induced transcriptional responses in hepatic cancer cells.

**Figure 7.**
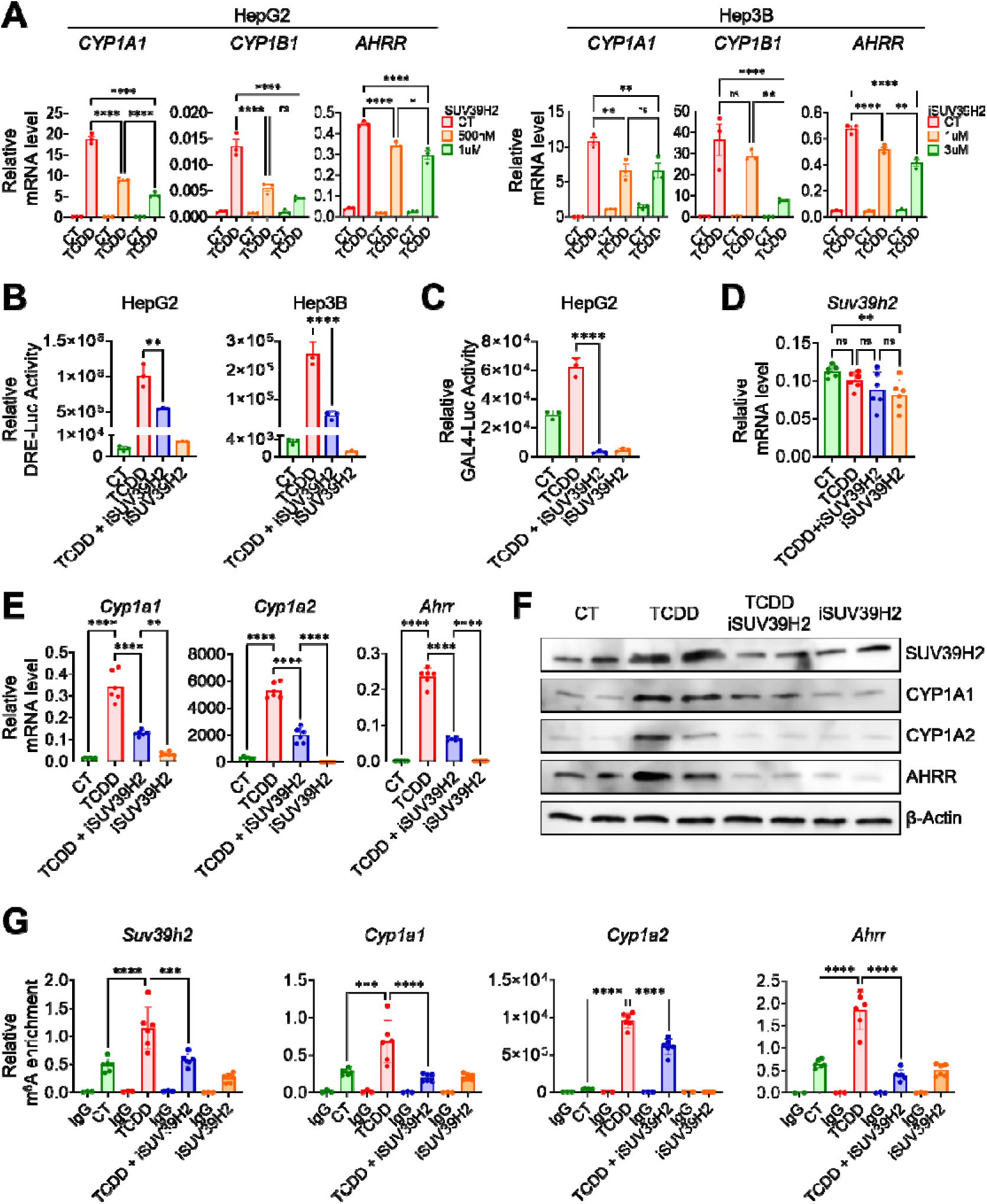
Pharmacological inhibition of SUV39H2 attenuates TCDD-induced AHR transcriptional responses in vitro and in vivo. **(A)** RT-qPCR analysis of *CYP1A1*, *CYP1B1*, and *AHRR* mRNA expression in HepG2 and Hep3B cells following TCDD exposure in the presence or absence of the SUV39H2 inhibitor OTS186935. HepG2 cells were treated with 500 nM or 1 µM inhibitor, whereas Hep3B cells were treated with 1 or 3 µM inhibitor. Transcript levels were normalized to *ACTB* expression. **(B)** Dioxin response element (DRE)-driven luciferase reporter activity in HepG2 and Hep3B cells treated with vehicle (CT), TCDD, TCDD plus OTS186935, or OTS186935 alone. **(C)** GAL4-dependent luciferase reporter activity in HepG2 cells following treatment with vehicle, TCDD, TCDD plus OTS186935, or OTS186935 alone. **(D)** RT-qPCR analysis of hepatic *Suv39h2* mRNA expression in mice treated with vehicle (CT), TCDD, TCDD plus OTS186935, or OTS186935 alone. **(E)** RT-qPCR analysis of hepatic *Cyp1a1*, *Cyp1a2*, and *Ahrr* mRNA expression in the indicated mouse treatment groups. Transcript levels were normalized to *Actb* expression. **(F)** Representative immunoblots showing hepatic SUV39H2, CYP1A1, CYP1A2, and AHRR protein abundance in the indicated mouse treatment groups, with β-actin serving as a loading control. **(G)** MeRIP-qPCR analysis of m⁶A enrichment in *Suv39h2*, *Cyp1a1*, *Cyp1a2*, and *Ahrr* transcripts in mouse liver following the indicated treatments. Enrichment was normalized to the corresponding input RNA, with IgG immunoprecipitation serving as a negative control. Statistical significance is indicated as *P* < 0.05 (*), *P* < 0.01 (**), *P* < 0.001 (***), and *P* < 0.0001 (****); ns, not significant. Statistical methods are described in Materials and Methods.

To determine whether these effects extend to an in vivo setting, we examined hepatic AHR-responsive gene expression in mice treated with TCDD, the SUV39H2 inhibitor, or their combination. Hepatic *Suv39h2* mRNA abundance remained largely unchanged following TCDD exposure or inhibitor treatment alone, whereas a modest reduction was observed following combined treatment (**Fig. 7D**). TCDD markedly induced hepatic *Cyp1a1*, *Cyp1a2*, and *Ahrr* mRNA expression, and these responses were significantly attenuated by SUV39H2 inhibitor co-treatment (**Fig. 7E**). Immunoblot analysis similarly showed reduced TCDD-induced CYP1A1, CYP1A2, and AHRR protein abundance following inhibitor co-treatment (**Fig. 7F**), supporting the involvement of SUV39H2 in regulating hepatic AHR transcriptional responses in vivo.

We next investigated whether pharmacological inhibition of SUV39H2 was associated with changes in m⁶A enrichment of AHR-responsive transcripts in mouse liver. MeRIP-qPCR analysis revealed increased m⁶A enrichment of *Suv39h2*, *Cyp1a1*, *Cyp1a2*, and *Ahrr* following TCDD exposure compared with vehicle-treated controls (**Fig. 7G**). Co-treatment with the SUV39H2 inhibitor attenuated TCDD-associated m⁶A enrichment of these transcripts, although the magnitude of reduction varied among individual genes. These observations indicate that SUV39H2 inhibitor treatment is associated with altered m⁶A enrichment of AHR-responsive transcripts in vivo, but do not establish whether SUV39H2 directly regulates RNA methylation.

Collectively, these findings demonstrate that pharmacological inhibition of SUV39H2 attenuates TCDD-induced AHR transcriptional responses in hepatic cancer cells and mouse liver. Together with the genetic loss- and gain-of-function experiments (**Fig. 6**), these results support a positive regulatory role for SUV39H2 in AHR signaling. The accompanying changes in transcript m⁶A enrichment further suggest a potential relationship between SUV39H2 activity and TCDD-associated RNA methylation, although the underlying mechanisms remain to be determined.

## Discussion

The aryl hydrocarbon receptor (AHR) mediates cellular responses to environmental chemicals through ligand-dependent transcriptional regulation. Although the canonical AHR signaling pathway has been extensively characterized, additional mechanisms controlling the magnitude and duration of AHR-dependent transcription remain incompletely understood (Denison and Nagy 2003; Beischlag et al. 2008; Safe et al. 2020; Kim 2024). In the present study, we investigated TCDD-associated changes in N⁶-methyladenosine (m⁶A) RNA modification and identified SUV39H2 as a previously unrecognized positive regulator of AHR signaling. TCDD increased m⁶A enrichment, apparent mRNA stability, and protein abundance of SUV39H2 without substantially altering its steady-state transcript levels. Importantly, SUV39H2 depletion and pharmacological inhibition attenuated TCDD-induced AHR target gene expression, whereas SUV39H2 overexpression enhanced AHR-dependent transcriptional responses. These findings suggest that SUV39H2 contributes to the regulation of AHR signaling and may provide a functional connection between post-transcriptional RNA regulation and chromatin-associated transcriptional control.

Exploratory MeRIP-seq analysis revealed TCDD-associated changes in m⁶A enrichment at selected transcripts despite largely unchanged global m⁶A abundance, methyltransferase activity, and transcriptome-wide distribution patterns. Consistent with previous studies, m⁶A enrichment was preferentially detected near the coding sequence–3′ untranslated region boundary (Dominissini et al. 2012; Meyer et al. 2012). Independent MeRIP-qPCR and mRNA decay analyses demonstrated increased m⁶A enrichment and prolonged apparent half-lives of canonical AHR-responsive transcripts, including *CYP1A1*, *CYP1A2*, and *AHRR*. These observations suggest that post-transcriptional regulation may contribute to the sustained expression of AHR target genes following TCDD exposure. Supporting this possibility, Shen et al. (2024) reported a role for m⁶A modification in regulating *CYP1A1* expression following naphtho[2,1-*a*]pyrene exposure, whereas Nakano et al. (2025) implicated m⁶A-associated regulatory mechanisms in *CYP1A1* induction involving ARNT and AHRR. However, the relationship between m⁶A enrichment and gene expression was heterogeneous across the transcripts examined. This observation is consistent with the diverse functions of m⁶A in RNA processing, stability, and translation, which depend on transcript context and the proteins recognizing modified RNA (Wang et al. 2014; Wang et al. 2015; Roundtree et al. 2017; Huang et al. 2018). Thus, TCDD-associated m⁶A remodeling may influence gene expression through transcript-specific mechanisms rather than uniformly promoting mRNA accumulation.

Among the candidate m⁶A-enriched transcripts, *SUV39H2* was particularly notable because of its established role in chromatin regulation. TCDD increased m⁶A enrichment and apparent mRNA stability of *SUV39H2*, accompanied by increased protein abundance despite minimal changes in steady-state transcript levels. These findings are consistent with post-transcriptional regulation of SUV39H2 following TCDD exposure. SUV39H2 is conventionally recognized as a histone H3 lysine 9 methyltransferase associated with repressive chromatin modifications. However, its transcriptional functions may be context dependent. Askew et al. (2017) demonstrated that SUV39H2 participates in androgen receptor-associated transcriptional regulation, providing a precedent for its involvement in nuclear receptor signaling beyond conventional transcriptional repression. Our findings extend this concept to AHR signaling. SUV39H2 depletion reduced TCDD-induced DRE-driven reporter activity and endogenous AHR target gene expression in HepG2 and Hep3B cells, whereas SUV39H2 overexpression enhanced these responses. Furthermore, CUT&RUN-qPCR revealed increased occupancy of both SUV39H2 and AHR at the *CYP1A1* enhancer following TCDD exposure. Because AHR-responsive regulatory elements recruit multiple transcriptional regulators during ligand-dependent activation (Beischlag et al. 2008; Matthews et al. 2005), the enrichment of SUV39H2 at this region supports its potential involvement in AHR-associated chromatin regulation. Together, these findings are consistent with a positive, coactivator-like function of SUV39H2 in AHR signaling, although direct interaction with AHR and the requirement for SUV39H2 catalytic activity remain to be established.

The positive regulatory role of SUV39H2 was further supported by pharmacological and *in vivo* experiments. OTS186935, a previously characterized small-molecule inhibitor of SUV39H2 methyltransferase activity, has been investigated for its effects on cancer-associated cellular processes (Vougiouklakis et al. 2018). In the present study, OTS186935 attenuated TCDD-induced AHR target gene expression and DRE-driven reporter activity in hepatic cancer cells, consistent with the genetic loss-of-function findings. Moreover, inhibitor co-treatment reduced TCDD-induced hepatic *Cyp1a1*, *Cyp1a2*, and *Ahrr* expression at both the mRNA and protein levels in mice. These observations suggest that SUV39H2 contributes to the regulation of AHR-responsive transcription beyond cultured cells. Interestingly, OTS186935 co-treatment also reduced TCDD-associated m⁶A enrichment of *Suv39h2*, *Cyp1a1*, *Cyp1a2*, and *Ahrr* transcripts in mouse liver. Because SUV39H2 is a histone methyltransferase rather than an established RNA m⁶A writer, these findings should not be interpreted as evidence that SUV39H2 directly catalyzes RNA methylation. Instead, they raise the possibility that SUV39H2-associated transcriptional regulation and RNA methylation responses are functionally interconnected. Whether the observed changes in m⁶A enrichment reflect altered RNA methylation machinery, AHR-dependent transcriptional responses, or indirect effects of inhibitor treatment requires further investigation. Given the established importance of AHR in xenobiotic metabolism and chemical-induced toxicity (Denison and Nagy 2003; Safe et al. 2013), SUV39H2-dependent regulation may have broader toxicological implications.

Based on these observations, we propose a working model linking TCDD-induced AHR activation, m⁶A-associated post-transcriptional regulation, and SUV39H2-dependent transcriptional responses (**Fig. 8**). In this model, TCDD activates AHR signaling and is accompanied by increased m⁶A enrichment of *SUV39H2* transcripts. Increased m⁶A enrichment coincides with prolonged apparent transcript stability and elevated SUV39H2 protein abundance, potentially increasing the availability of SUV39H2 for transcriptional regulatory functions. SUV39H2 may subsequently enhance AHR-dependent transcription through its association with AHR-responsive chromatin, thereby contributing to the induction of downstream target genes, including *CYP1A1*, *CYP1A2*, and *AHRR*. This model integrates the post-transcriptional and transcriptional observations into a potential regulatory pathway. However, the proposed connections between m⁶A modification, SUV39H2 protein accumulation, and chromatin-associated AHR activation remain mechanistic hypotheses rather than established causal relationships.

**Figure 8.**
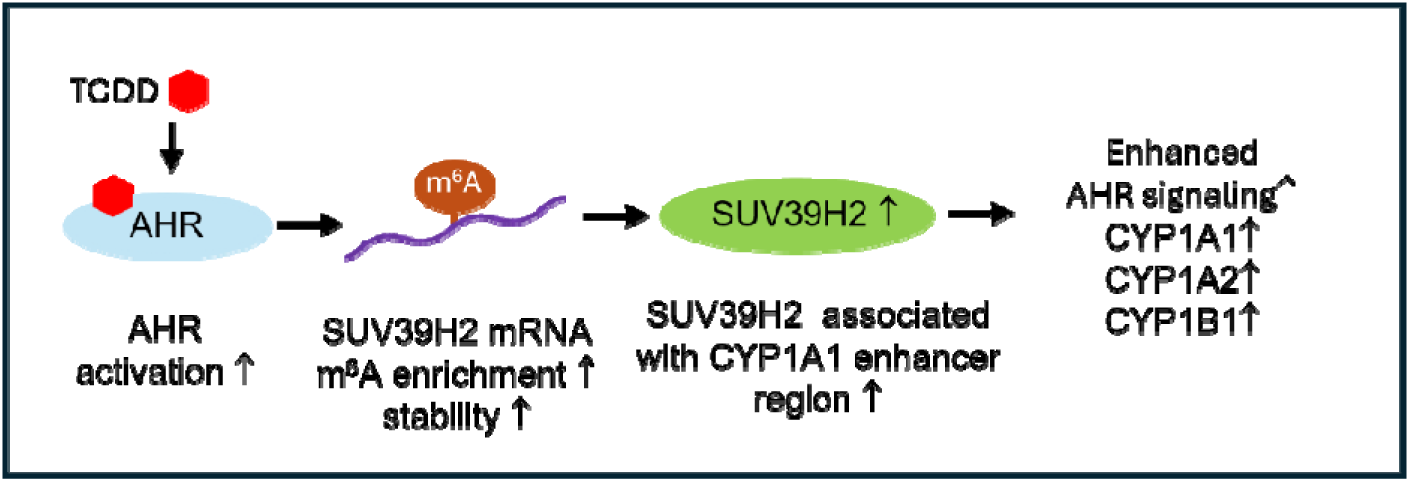
Proposed model linking TCDD-induced AHR activation, m⁶A-associated post-transcriptional regulation, and SUV39H2 expression. TCDD activates AHR signaling and is associated with increased m⁶A enrichment of *SUV39H2* transcripts. This modification coincides with increased apparent mRNA stability and elevated SUV39H2 protein abundance. The proposed model suggests that increased SUV39H2 availability may contribute to enhanced AHR-dependent transcriptional responses. The functional relationship between m⁶A enrichment and SUV39H2 stabilization, as well as the molecular mechanism underlying SUV39H2-mediated regulation of AHR activity, remains to be established.

Several limitations should be acknowledged. First, MeRIP-seq and matched RNA-seq were performed using one biological sample per condition; therefore, transcriptome-wide findings were considered exploratory, and conclusions regarding selected transcripts were supported by independent validation experiments. Second, although increased m⁶A enrichment was associated with prolonged apparent *SUV39H2* mRNA stability and elevated protein abundance, a direct causal relationship has not been demonstrated. Identification of individual m⁶A sites, characterization of relevant RNA-binding proteins, and perturbation of the RNA methylation machinery would help clarify whether m⁶A directly contributes to SUV39H2 stabilization or translation. Third, whether SUV39H2 directly interacts with AHR or regulates AHR-dependent transcription through its histone methyltransferase activity remains unknown. Protein-interaction studies, catalytic-mutant rescue experiments, and chromatin analyses would help distinguish these mechanisms. Furthermore, although genetic and pharmacological approaches produced consistent effects, potential off-target activities of OTS186935 cannot be excluded. Finally, the *in vivo* experiments primarily examined short-term hepatic transcriptional responses rather than downstream toxicological outcomes, such as changes in xenobiotic metabolism, inflammation, or liver injury.

In conclusion, this study identifies SUV39H2 as a previously unrecognized positive regulator of TCDD-induced AHR signaling. TCDD-associated increases in *SUV39H2* m⁶A enrichment, apparent transcript stability, and protein abundance, together with complementary genetic, pharmacological, and *in vivo* findings, support a potential functional connection between RNA methylation-associated post-transcriptional regulation and chromatin-associated AHR transcriptional responses. The proposed model (**Fig. 8**) provides a framework for investigating how environmental AHR ligands may influence transcriptional responses through interconnected epitranscriptomic and epigenetic mechanisms. These findings expand our understanding of AHR signaling and identify SUV39H2 as a candidate regulator of cellular responses to environmental chemicals.

## Author Contributions

Conceptualization, K. Kim; Methodology, H.D. Yun, G. Kwon, V. Wanchai, and K. Kim; Validation, H.D. Yun, G. Kwon, and K. Kim; Formal analysis, all authors; Investigation, K. Kim, H.D. Yun, G. Kwon, and V. Wanchai; Resources, all authors; Data curation, K. Kim; Writing—original draft preparation, K. Kim, G. Kwon, and H.D. Yun; Writing—review and editing, K. Kim, G. Kwon, and S. Safe; Supervision, K. Kim; Project administration, K. Kim; Funding acquisition, K. Kim. All authors have read and approved the final version of the manuscript.

## Abbreviations

The following abbreviations are used in this manuscript:

AHR: aryl hydrocarbon receptor
AHRR: aryl hydrocarbon receptor repressor
AR: androgen eptor
ARNT: aryl hydrocarbon receptor nuclear translocator
ActD: actinomycin D
BCA: bicinchoninic acid
CDS: coding sequence
ChIP-qPCR: chromatin immunoprecipitation followed by quantitative polymerase chain reaction
CUT&RUN-qPCR: cleavage under targets and release using nuclease followed by quantitative polymerase chain reaction
CYP: cytochrome P450
DMSO: dimethyl sulfoxide
DRE: dioxin response element
FDR: false discovery rate
FPKM: fragments per kilobase of transcript per million mapped reads
H3K9: histone H3 lysine 9
IGF2BP: insulin-like growth factor 2 mRNA-binding protein
IP: immunoprecipitation
KEGG: Kyoto Encyclopedia of Genes and Genomes
m⁶A: N⁶-methyladenosine
MeRIP-qPCR: methylated RNA immunoprecipitation followed by quantitative polymerase chain reaction
MeRIP-seq: methylated RNA immunoprecipitation sequencing
METTL3: methyltransferase-like protein 3
METTL14: methyltransferase-like protein 14
RT-qPCR: reverse transcription quantitative polymerase chain reaction
SD: standard deviation
siRNA: small interfering RNA
SMN1: survival motor neuron 1
SUV39H2: suppressor of variegation 3-9 homolog 2
TCDD: 2,3,7,8-tetrachlorodibenzo-*p*-dioxin
UTR: untranslated region.

