## Supplementary material for "TCDD-Induced m⁶A Profiling Identifies SUV39H2 as a Positive Regulator of Aryl Hydrocarbon Receptor Signaling": Supplemetary information

### Slide 1
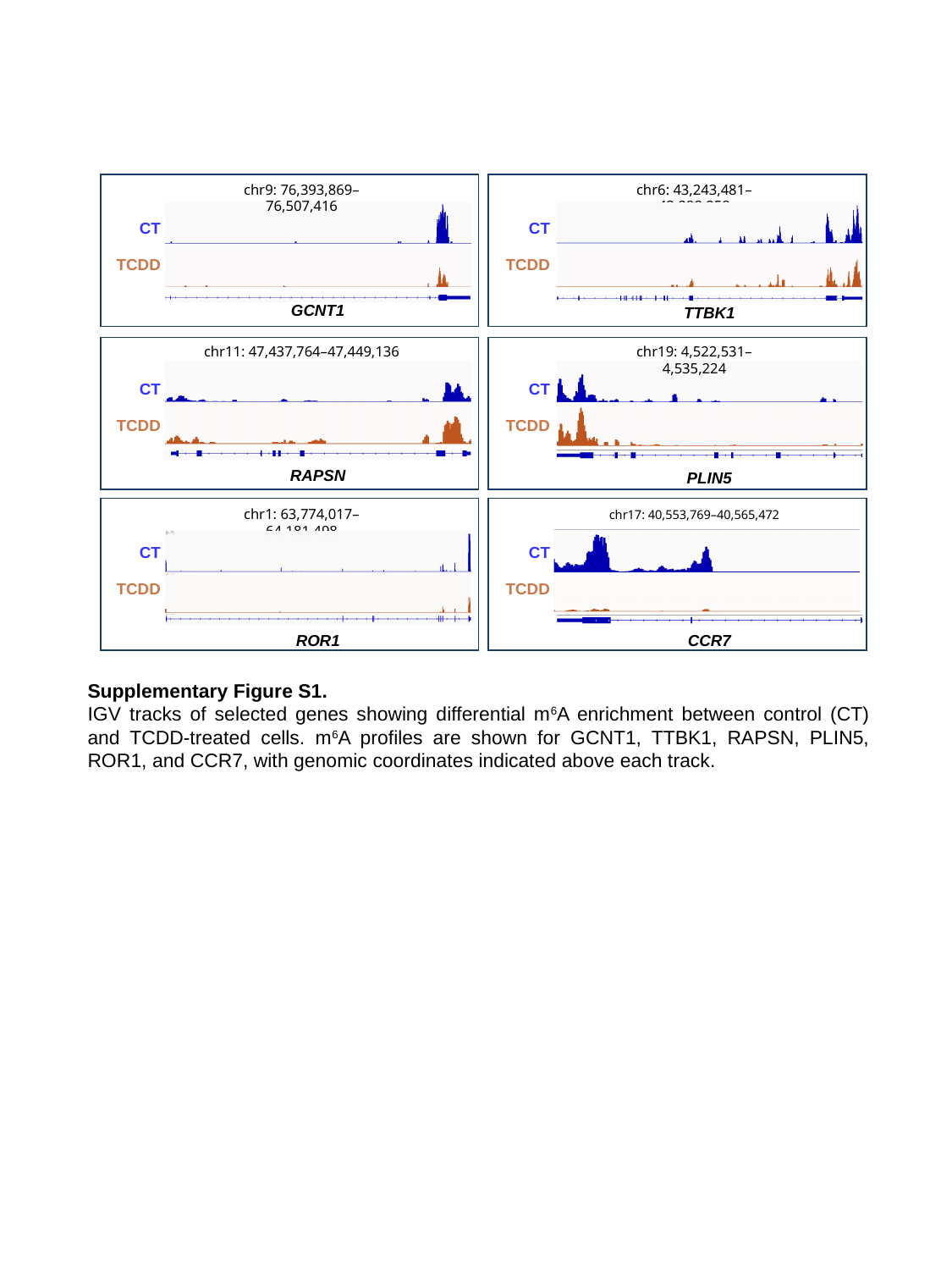

chr9: 76,393,869–76,507,416
chr6: 43,243,481–43,288,258
CT
CT
TCDD
TCDD
GCNT1
TTBK1
chr11: 47,437,764–47,449,136
chr19: 4,522,531–4,535,224
CT
CT
TCDD
TCDD
RAPSN
PLIN5
chr1: 63,774,017–64,181,498
chr17: 40,553,769–40,565,472
CT
CT
TCDD
TCDD
ROR1
CCR7
Supplementary Figure S1.
IGV tracks of selected genes showing differential m^6^A enrichment between control (CT) and TCDD-treated cells. m^6^A profiles are shown for GCNT1, TTBK1, RAPSN, PLIN5, ROR1, and CCR7, with genomic coordinates indicated above each track.

### Slide 2
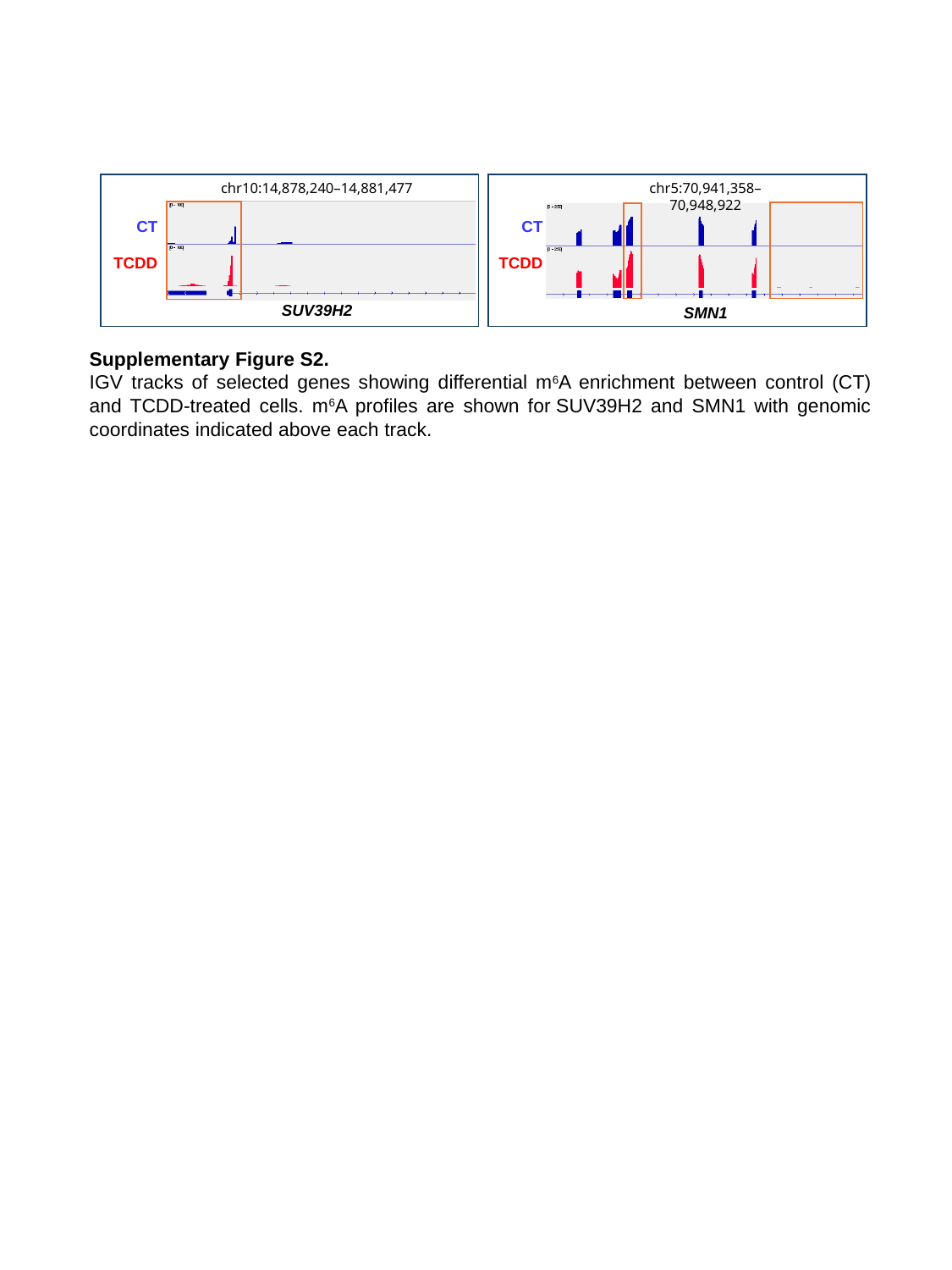

chr10:14,878,240–14,881,477
chr5:70,941,358–70,948,922
CT
CT
TCDD
TCDD
SUV39H2
SMN1
Supplementary Figure S2.
IGV tracks of selected genes showing differential m^6^A enrichment between control (CT) and TCDD-treated cells. m^6^A profiles are shown for SUV39H2 and SMN1 with genomic coordinates indicated above each track.

### Slide 3
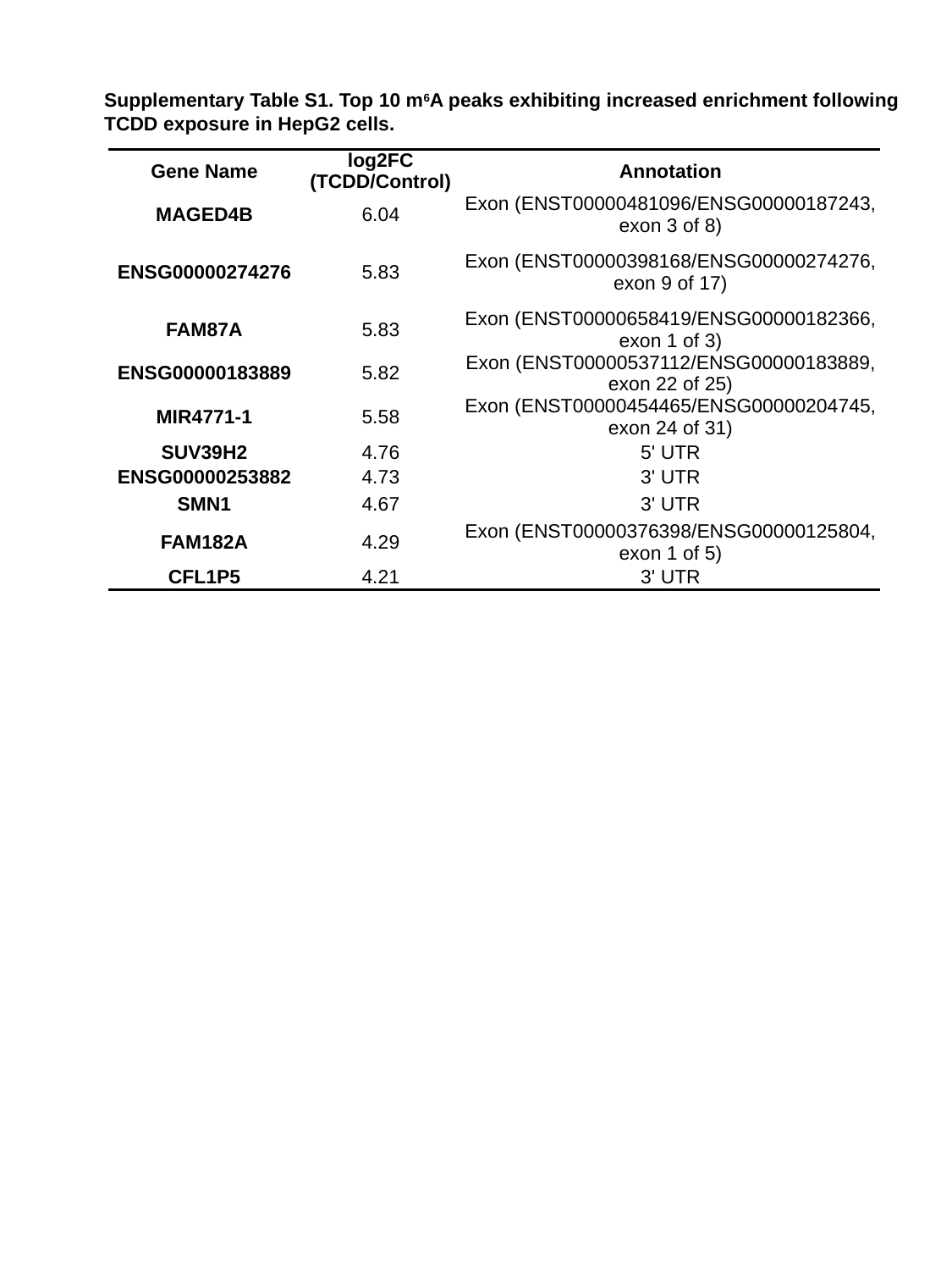

Supplementary Table S1. Top 10 m^6^A peaks exhibiting increased enrichment following TCDD exposure in HepG2 cells.
| Gene Name | log2FC (TCDD/Control) | Annotation |
| --- | --- | --- |
| MAGED4B | 6.04 | Exon (ENST00000481096/ENSG00000187243, exon 3 of 8) |
| ENSG00000274276 | 5.83 | Exon (ENST00000398168/ENSG00000274276, exon 9 of 17) |
| FAM87A | 5.83 | Exon (ENST00000658419/ENSG00000182366, exon 1 of 3) |
| ENSG00000183889 | 5.82 | Exon (ENST00000537112/ENSG00000183889, exon 22 of 25) |
| MIR4771-1 | 5.58 | Exon (ENST00000454465/ENSG00000204745, exon 24 of 31) |
| SUV39H2 | 4.76 | 5' UTR |
| ENSG00000253882 | 4.73 | 3' UTR |
| SMN1 | 4.67 | 3' UTR |
| FAM182A | 4.29 | Exon (ENST00000376398/ENSG00000125804, exon 1 of 5) |
| CFL1P5 | 4.21 | 3' UTR |

### Slide 4
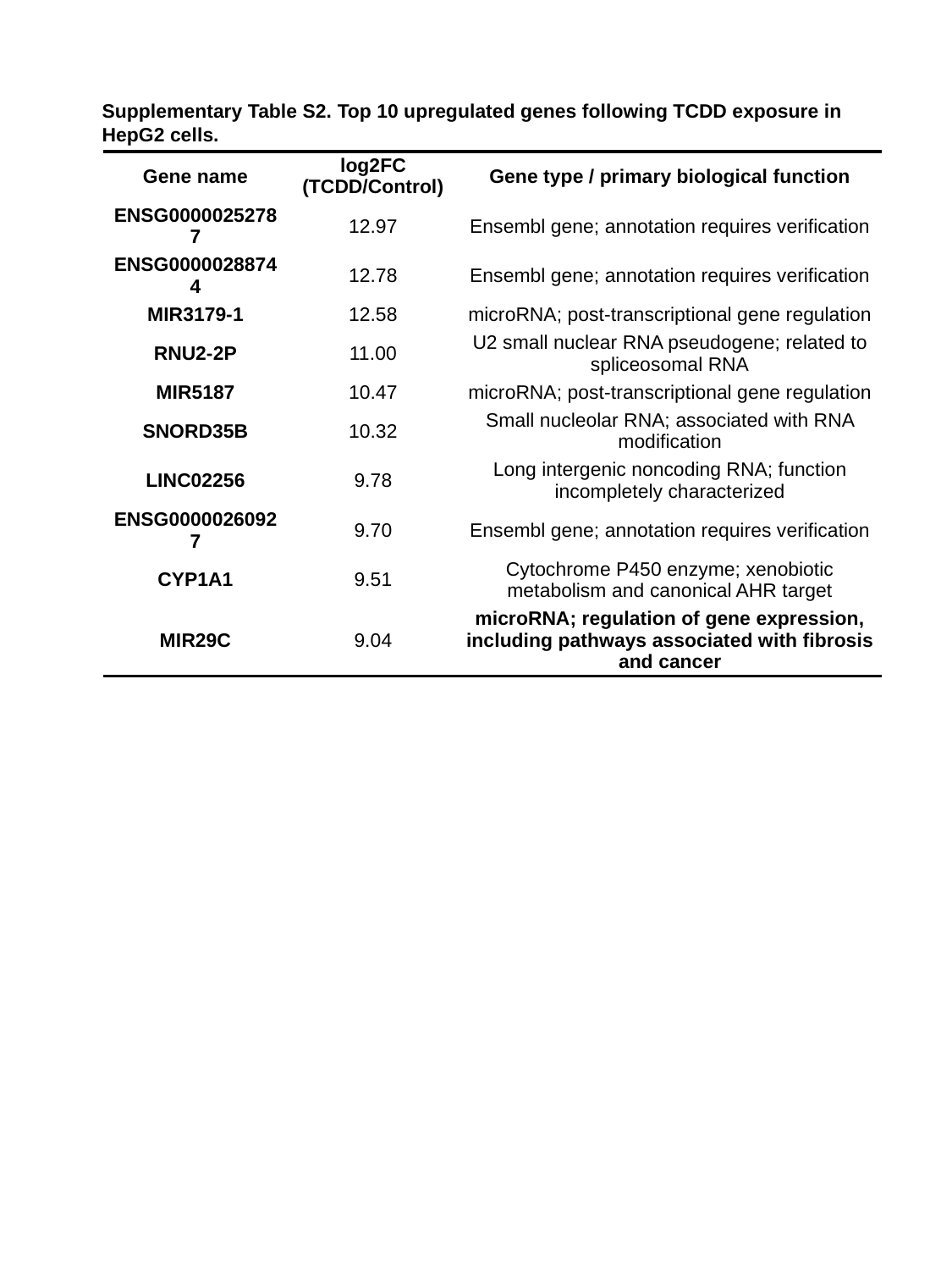

Supplementary Table S2. Top 10 upregulated genes following TCDD exposure in HepG2 cells.
| Gene name | log2FC (TCDD/Control) | Gene type / primary biological function |
| --- | --- | --- |
| ENSG00000252787 | 12.97 | Ensembl gene; annotation requires verification |
| ENSG00000288744 | 12.78 | Ensembl gene; annotation requires verification |
| MIR3179-1 | 12.58 | microRNA; post-transcriptional gene regulation |
| RNU2-2P | 11.00 | U2 small nuclear RNA pseudogene; related to spliceosomal RNA |
| MIR5187 | 10.47 | microRNA; post-transcriptional gene regulation |
| SNORD35B | 10.32 | Small nucleolar RNA; associated with RNA modification |
| LINC02256 | 9.78 | Long intergenic noncoding RNA; function incompletely characterized |
| ENSG00000260927 | 9.70 | Ensembl gene; annotation requires verification |
| CYP1A1 | 9.51 | Cytochrome P450 enzyme; xenobiotic metabolism and canonical AHR target |
| MIR29C | 9.04 | microRNA; regulation of gene expression, including pathways associated with fibrosis and cancer |

### Slide 5
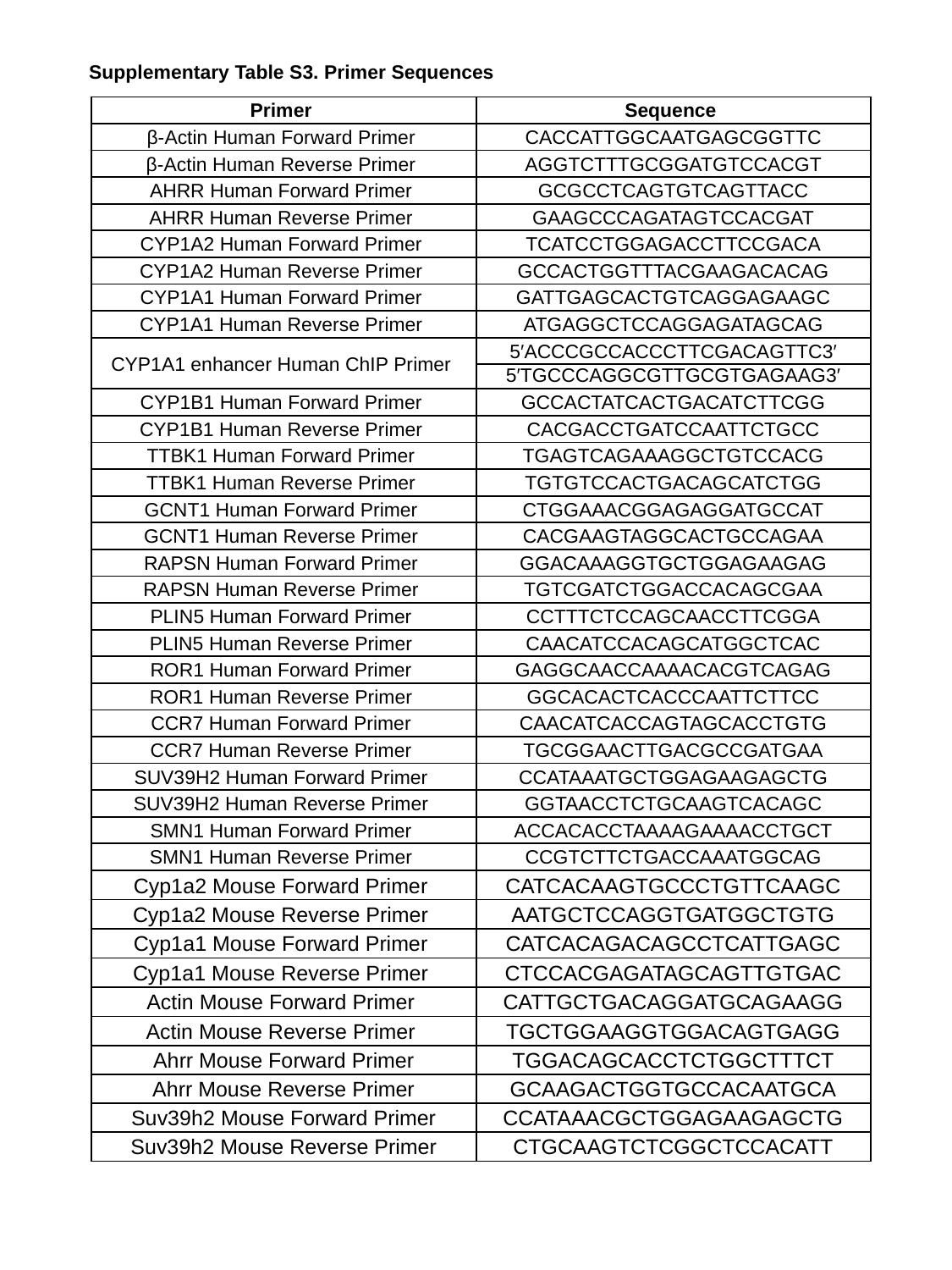

Supplementary Table S3. Primer Sequences
| Primer | Sequence |
| --- | --- |
| β-Actin Human Forward Primer | CACCATTGGCAATGAGCGGTTC |
| β-Actin Human Reverse Primer | AGGTCTTTGCGGATGTCCACGT |
| AHRR Human Forward Primer | GCGCCTCAGTGTCAGTTACC |
| AHRR Human Reverse Primer | GAAGCCCAGATAGTCCACGAT |
| CYP1A2 Human Forward Primer | TCATCCTGGAGACCTTCCGACA |
| CYP1A2 Human Reverse Primer | GCCACTGGTTTACGAAGACACAG |
| CYP1A1 Human Forward Primer | GATTGAGCACTGTCAGGAGAAGC |
| CYP1A1 Human Reverse Primer | ATGAGGCTCCAGGAGATAGCAG |
| CYP1A1 enhancer Human ChIP Primer | 5′ACCCGCCACCCTTCGACAGTTC3′ |
| | 5′TGCCCAGGCGTTGCGTGAGAAG3′ |
| CYP1B1 Human Forward Primer | GCCACTATCACTGACATCTTCGG |
| CYP1B1 Human Reverse Primer | CACGACCTGATCCAATTCTGCC |
| TTBK1 Human Forward Primer | TGAGTCAGAAAGGCTGTCCACG |
| TTBK1 Human Reverse Primer | TGTGTCCACTGACAGCATCTGG |
| GCNT1 Human Forward Primer | CTGGAAACGGAGAGGATGCCAT |
| GCNT1 Human Reverse Primer | CACGAAGTAGGCACTGCCAGAA |
| RAPSN Human Forward Primer | GGACAAAGGTGCTGGAGAAGAG |
| RAPSN Human Reverse Primer | TGTCGATCTGGACCACAGCGAA |
| PLIN5 Human Forward Primer | CCTTTCTCCAGCAACCTTCGGA |
| PLIN5 Human Reverse Primer | CAACATCCACAGCATGGCTCAC |
| ROR1 Human Forward Primer | GAGGCAACCAAAACACGTCAGAG |
| ROR1 Human Reverse Primer | GGCACACTCACCCAATTCTTCC |
| CCR7 Human Forward Primer | CAACATCACCAGTAGCACCTGTG |
| CCR7 Human Reverse Primer | TGCGGAACTTGACGCCGATGAA |
| SUV39H2 Human Forward Primer | CCATAAATGCTGGAGAAGAGCTG |
| SUV39H2 Human Reverse Primer | GGTAACCTCTGCAAGTCACAGC |
| SMN1 Human Forward Primer | ACCACACCTAAAAGAAAACCTGCT |
| SMN1 Human Reverse Primer | CCGTCTTCTGACCAAATGGCAG |
| Cyp1a2 Mouse Forward Primer | CATCACAAGTGCCCTGTTCAAGC |
| Cyp1a2 Mouse Reverse Primer | AATGCTCCAGGTGATGGCTGTG |
| Cyp1a1 Mouse Forward Primer | CATCACAGACAGCCTCATTGAGC |
| Cyp1a1 Mouse Reverse Primer | CTCCACGAGATAGCAGTTGTGAC |
| Actin Mouse Forward Primer | CATTGCTGACAGGATGCAGAAGG |
| Actin Mouse Reverse Primer | TGCTGGAAGGTGGACAGTGAGG |
| Ahrr Mouse Forward Primer | TGGACAGCACCTCTGGCTTTCT |
| Ahrr Mouse Reverse Primer | GCAAGACTGGTGCCACAATGCA |
| Suv39h2 Mouse Forward Primer | CCATAAACGCTGGAGAAGAGCTG |
| Suv39h2 Mouse Reverse Primer | CTGCAAGTCTCGGCTCCACATT |

### Slide 6
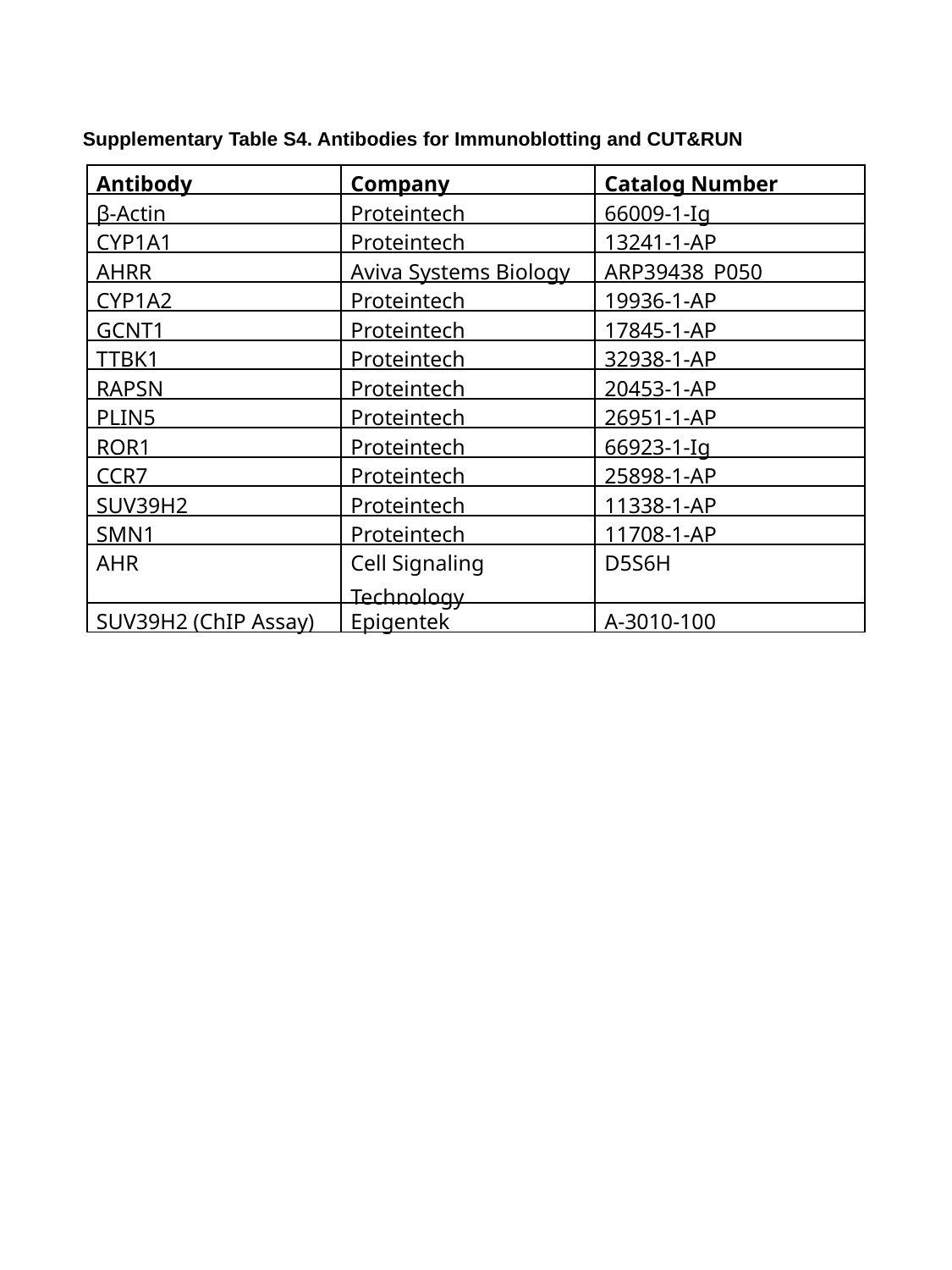

Supplementary Table S4. Antibodies for Immunoblotting and CUT&RUN
| Antibody | Company | Catalog Number |
| --- | --- | --- |
| β-Actin | Proteintech | 66009-1-Ig |
| CYP1A1 | Proteintech | 13241-1-AP |
| AHRR | Aviva Systems Biology | ARP39438\_P050 |
| CYP1A2 | Proteintech | 19936-1-AP |
| GCNT1 | Proteintech | 17845-1-AP |
| TTBK1 | Proteintech | 32938-1-AP |
| RAPSN | Proteintech | 20453-1-AP |
| PLIN5 | Proteintech | 26951-1-AP |
| ROR1 | Proteintech | 66923-1-Ig |
| CCR7 | Proteintech | 25898-1-AP |
| SUV39H2 | Proteintech | 11338-1-AP |
| SMN1 | Proteintech | 11708-1-AP |
| AHR | Cell Signaling Technology | D5S6H |
| SUV39H2 (ChIP Assay) | Epigentek | A-3010-100 |
